# Molecular fingerprints for a novel glucosamine kinase family in *Actinobacteria*

**DOI:** 10.1101/482810

**Authors:** José A Manso, Daniela Nunes-Costa, Sandra Macedo-Ribeiro, Nuno Empadinhas, Pedro José Barbosa Pereira

## Abstract

*Actinobacteria* have long been the main source of antibiotics, secondary metabolites with tightly controlled biosynthesis by environmental and physiological factors. Phosphorylation of exogenous glucosamine has been suggested as a mechanism for incorporation of this extracellular material into secondary metabolite biosynthesis, but experimental evidence of specific glucosamine kinases in *Actinobacteria* is lacking. Here, we present the molecular fingerprints for the identification of a unique family of actinobacterial glucosamine kinases. Structural and biochemical studies on a distinctive kinase from the soil bacterium *Streptacidiphilus jiangxiensis* unveiled its preference for glucosamine and provided structural evidence of a phosphoryl transfer to this substrate. Conservation of glucosamine-contacting residues across a large number of uncharacterized actinobacterial proteins unveiled a specific glucosamine-binding sequence motif. This family of kinases and their genetic context may represent the missing link for the incorporation of environmental glucosamine into the antibiotic biosynthesis pathways in *Actinobacteria* and can be explored to enhance antibiotic production.

## Introduction

The worrying worldwide escalation of antimicrobial resistance is limiting the treatment options for numerous infectious diseases (Fair and Tor, 2014). Therefore, finding new strategies that lead to either novel effective antibiotics or enhancement of the production of the known drugs stand as global challenges of utmost importance (Brown and Wright, 2016).

*Actinobacteria* have long been the largest natural source of antibiotics (Genilloud, 2017). Two-thirds of the currently known antibiotics are produced by organisms of the genus *Streptomyces*, the largest (>800 valid species) in the phylum *Actinobacteria* and a paradigm of secondary metabolite-producing microorganisms (Hopwood, 1999). *Streptomyces* spp. belong to the *Streptomycetaceae* family, which so far includes only three additional genera, *Allostreptomyces*, *Kitasatospora* and *Streptacidiphilus*, all very difficult to differentiate both by genotypic and phenotypic characteristics (Kämpfer et al., 2014; Huang et al., 2017).

Post-genomic era advances in the regulation of antibiotic production in *Actinobacteria* resulted from the identification of one of the most important regulators of N-acetylglucosamine (GlcNAc) uptake and metabolism and of degradation of chitin to GlcNAc, DasR from the GntR family (Colson et al., 2007; van der Heul et al., 2018; Rigali et al., 2006; Świątek-Połatyńska et al., 2015). GlcNAc is an important antibiotic elicitor, participating in signaling pathways leading to secondary metabolite production in *Streptomyces* (Ochi and Hosaka, 2013; Rigali et al., 2008; Zhu et al., 2014) and providing the glycosyl moieties of some antibiotics (Kudo et al., 2005).

A central molecule of GlcNAc metabolism with a crucial role in cell wall synthesis, glycolysis and nitrogen metabolism is glucosamine-6-phosphate (GlcN-6P) (Jolly et al., 1997; Mengin-Lecreulx and Heijenoort, 1994; Mengin-Lecreulx and van Heijenoort, 1996; Świątek et al., 2012). Moreover and most importantly, GlcN-6P binds to DasR and modulates its DNA-binding activity, regulating the activation of antibiotic production in Actinobacteria (Abdelmohsen et al., 2015; Ochi and Hosaka, 2013; Rigali et al., 2008; Zhu et al., 2014). The crystal structures of DasR and of its ortholog NagR from *Bacillus subtilis* in complex with GlcN-6P have been reported (Fillenberg et al., 2015, 2016). However, to date two metabolic enzymes that lead to production of intracellular GlcN-6P in *Actinobacteria* have been described: N-acetylglucosamine-6-phosphate (GlcNAc-6P) deacetylase, NagA (EC 3.5.1.25), that deacetylates GlcNAc-6P formed by phosphorylation of acquired extracellular GlcNAc by a phosphotransferase system (Plumbridge and Vimr, 1999; Vogler and Lengeler, 1989; Plumbridge, 2009; Ahangar et al., 2018), and GlcN-6P synthase (EC 2.6.1.16), that produces GlcN-6P by transamination of the glycolytic intermediate fructose-6P (Milewski, 2002; Teplyakov et al., 2002). Homologues of these two enzyme classes are present in the genomes of several *Streptomycetaceae*. On the other hand, it was recently demonstrated that the *csnR-K* operon, conserved in *Actinobacteria*, controls the uptake of chitosan-derived D-glucosamine (GlcN) oligosaccharides and, in lower scale, of monomeric GlcN (Viens et al., 2015). Phosphorylation of exogenous GlcN to GlcN-6P could thus constitute an alternative mechanism for incorporation of extracellular material (chitin or chitosan) into the actinobacterial GlcN metabolism. Although GlcN kinases (EC 2.7.1.8) have been identified in the γ-proteobacterium *Vibrio cholera* (Park et al., 2002) and in the hyperthermophilic archaeon *Thermococcus kodakarensis* (Aslam et al., 2018), and their existence in *Actinobacteria* has also been anticipated, allowing incorporation of extracellular material (chitin or chitosan) into the GlcN metabolism, experimental evidence for this activity in that medically important phylum was still missing. Here, we identify and characterize a unique GlcN kinase from the soil bacterium *Streptacidiphilus jiangxiensis*, closely related to *Streptomyces* spp., the classical antibiotic producers (Huang et al., 2004). This GlcN kinase is not a sequence homologue of the isofunctional enzymes from *Vibrio* or from archaeal (Park et al., 2002; Aslam et al., 2018). The crystal structure of the quaternary complex between this unusual kinase and GlcN, ADP, inorganic phosphate and Mg^2+^, provides a unique structural evidence of a transition state of the phosphoryl-transfer mechanism in this unique family of GlcN kinases with eukaryotic protein kinase fold. In addition, the molecular determinants for GlcN phosphorylation, with high evolutive conservation in several representative families of the phylum *Actinobacteria* have been determined, unveiling a sequence motif for the classification of a large number of uncharacterized or misannotated aminoglycoside phosphotransferases into this unique family of actinobacterial GlcN kinases.

## Results

### A maltokinase paralogue associated with biosynthetic gene clusters in Actinobacteria

In a few actinobacterial genomes a gene with sequence homology to maltokinases (and annotated as such) was identified, in addition to the canonical maltokinase gene (Mendes et al., 2010). In each species, the sequence identity between the canonical maltokinase and its paralogue was around 30%. Maltokinases catalyze the synthesis of maltose-1-phosphate, a key metabolite in α-glucan biosynthesis in actinomycetes (Rashid et al., 2016). The presence of a putative second maltokinase gene in some actinobacteria prompted a careful analysis of its genomic context, which revealed a tight association with two other uncharacterized genes, a putative sugar isomerase (SIS) of the AgaS superfamily (COG2222) and a putative glycosyltransferase (GT) with homology to members of the GT1 family (Figure 1). In *Streptacidiphilus jiangxiensis*, two other genes with possible sugar-modifying activities are present, annotated as putative phosphoglucomutase (PGM) and UTP--glucose-1-phosphate uridylyltransferase (UDPGP), thus completing a potential pathway for sugar activation and transfer. Furthermore, in several actinomycetes, this operon was predicted to be included in larger biosynthetic gene clusters (BGCs), suggesting a role in the production of complex secondary metabolites (Figure 1). Since the *Streptacidiphilus jiangxiensis* homologue was part of an apparently more complete operon and given the phylogenetic relationship of this organism with antibiotic-producing streptomycetes, this gene was selected for further investigation.

**Figure 1.**
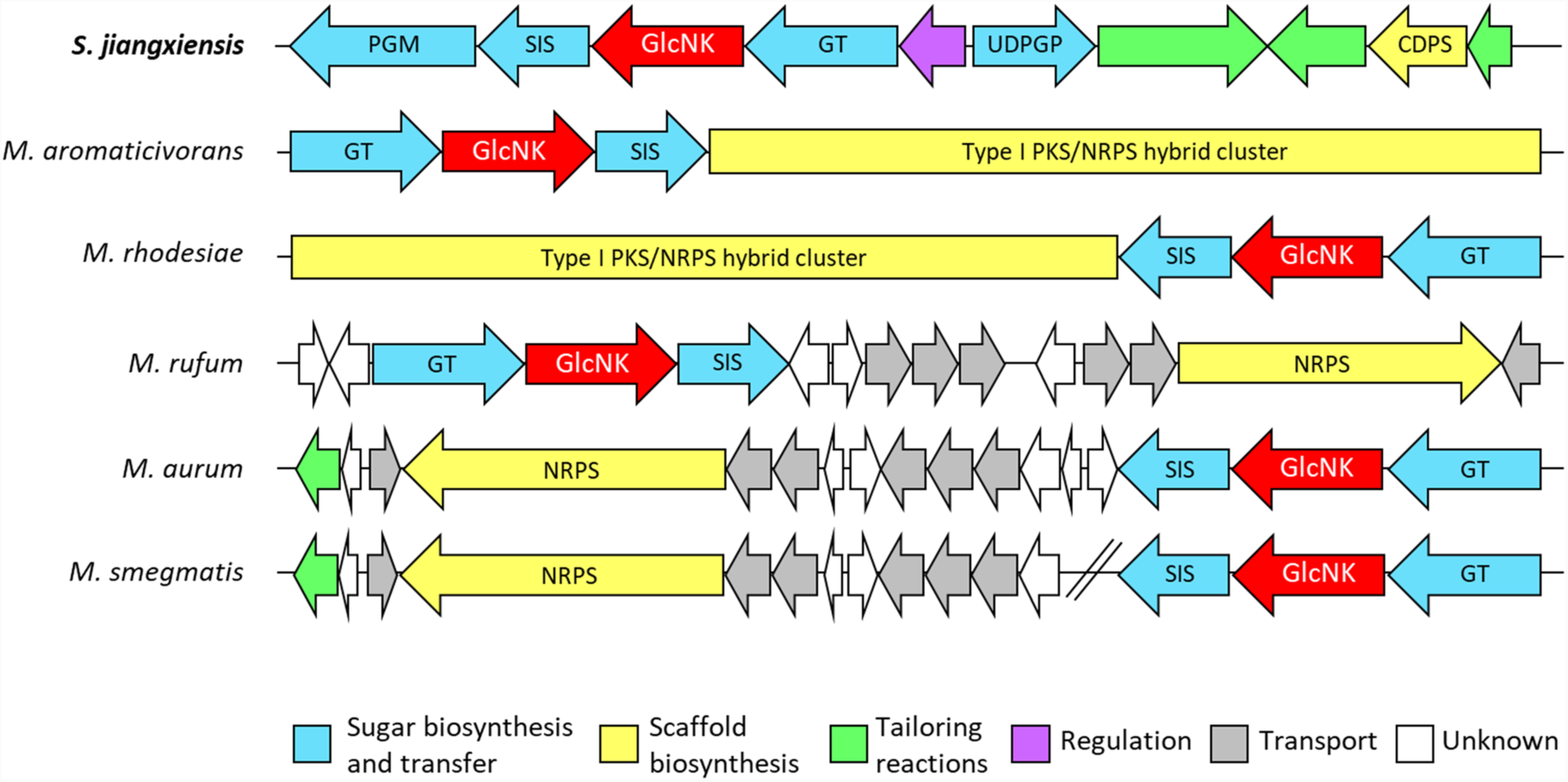
Genomic context of the *Streptacidiphilus jiangxiensis* maltokinase paralogue, SjGlcNK, and of its homologues in selected *Actinobacteria*. GlcNK is predicted to be within the borders of biosynthetic gene clusters in all represented species except for *Mycolicibacterium smegmatis*, which appears to have a cluster homologous to *Mycolicibacterium rufum* and *Mycolicibacterium aurum* but with the sugar-modifying genes, including GlcNK, located elsewhere on the chromosome. Figure drawn according to antiSMASH (Blin et al., 2017) results. PGM, putative phosphoglucomutase; SIS, putative phosphosugar isomerase; GlcNK, GlcN kinase; GT, putative glycosyltransferase; UDPGP, putative UTP--glucose-1-phosphate uridylyltransferase; CDPS, putative cyclodipeptide synthase; NRPS, putative nonribosomal peptide synthase; PKS, putative polyketide synthase.

### The S. jiangxiensis maltokinase paralogue encodes an enzyme with GlcN kinase activity

In order to gain more insight into the potential function of the *S. jiangxiensis* maltokinase paralogue, hereafter SjGlcNK, a sequence profile-based search for homologs was performed using the Fold and Function Assignment System, FFAS03 (Jaroszewski et al., 2011). As expected, three significant hits were identified corresponding to the maltokinases from *Mycolicibacterium smegmatis* (MsMak; PDB entry 5JY7, FFAS-score=−98.8), *Mycolicibacterium vanbaalenii* (MvMak, PDB entry 4U94 (Fraga et al., 2015), FFAS-score=−98.3) and *Mycobacterium tuberculosis* (MtMak, PDB entry 4O7O (Li et al., 2014), FFAS-score=−97.5). FFAS03 scores below −9.5 correspond to high confidence predictions with less than 3% false positives. The mycobacterial homologs identified display 21%-22% amino acid sequence identity to SjGlcNK, with many of the residues of the canonical structural motifs associated with nucleotide binding and enzymatic activity in the maltokinases being highly conserved in SjGlcNK (Figure S1). Despite a slightly shorter sequence, the important catalytic motifs P-loop (^117^VDQTNESV^124^), the ^130^AVVKW^134^ region containing the conserved phosphate-binding lysine residue, as well as the catalytic (^296^DVHGDFHVGQI^306^) and ^317^DFD^319^ loops could be identified in SjGlcNK (Figure S1, Figure 2*A*). Interestingly, SjGlcNK was unable to phosphorylate maltose *in vitro* (Figure 2*B*) indicating an incorrect annotation of its function. In contrast, from 20 sugars and 4 aminoglycoside antibiotics tested, SjGlcNK only phosphorylated specifically GlcN and displayed vestigial activity with glucose using ATP as phosphate donor (Figure 2*C*). GlcN was modified at position 6 yielding GlcN-6P, as determined by NMR (Figure 2*D*, Figure S2). Determination of the Michaelis constants (*K*_m_) for glucose and GlcN suggested that SjGlcNK is better described as a GlcN kinase (EC 2.7.1.8) with a strong preference for using GlcN (*K*_m_ = 8 ± 1 mM) over glucose (*K*_m_ > 100 mM) as substrate (Figure 2*E*).

**Figure 2.**
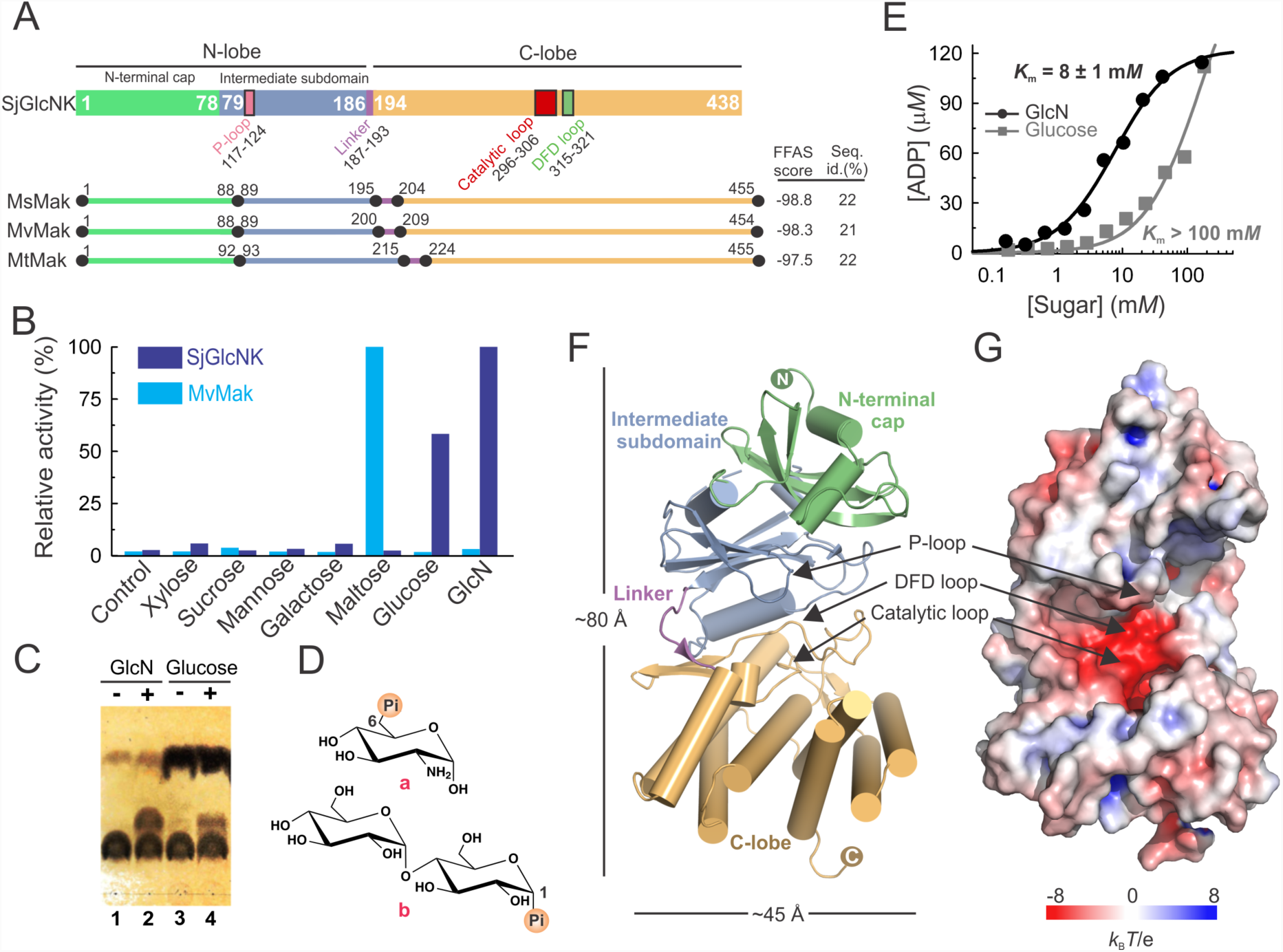
Biochemical characterization and overall structure of SjGlcNK. (*A*) Structural organization of SjGlcNK (colored rectangles) and of the mycobacterial maltokinases MsMak (PDB entry 5JY7), MvMak (PDB entry 4U94 (Fraga et al., 2015)) and MtMak (PDB entry 4O7O (Li et al., 2014)) (colored lines), indicating the relative arrangement of the N-terminal cap (light green) and intermediate subdomains (blue), linker (magenta) and C-lobe (orange). The canonical structural motifs in SjGlcNK-P-loop (pink), catalytic loop (red) and DFD loop (green) - are also indicated. The FFAS score (Jaroszewski et al., 2011) and the percentage of amino acid sequence identity between SjGlcNK and each of the mycobacterial maltokinases are given to the right. (*B*) Relative kinase activity of SjGlcNK (marine) and MvMak (cyan) towards different potential phosphate acceptors (xylose, sucrose, mannose, galactose, maltose, glucose and GlcN). The reaction mix ([enzyme] = 2 μ*M*, [sugar] = 5 m*M*, [ATP] = 1 m*M*) was incubated at RT for 5 min and activity was quantified with the ADP-Glo^™^ kinase assay. The results were normalized against the luminescence signal for the reactions with maltose (MvMak) or GlcN (SjGlcNK). While MvMak uses exclusively maltose as substrate acceptor, SjGlcNK phosphorylates glucose and GlcN, with preference for the latter. As control, the activity in the absence of phosphate-accepting sugars is shown. (*C*) Thin-layer chromatography analysis of SjGlcNK-catalyzed phosphorylation of GlcN and glucose using ATP as phosphate donor. Lanes 1 and 3: Negative controls without SjGlcNK. Lane 2: Reaction with GlcN. Lane 4: Reaction with glucose. (*D*) Chemical structures of the reaction products of SjGlcNK and MvMak, GlcN-6P (a) and maltose-1-phosphate (b), respectively. (*E*) Kinetics of SjGlcNK activity towards glucose and GlcN. The experimental data were fitted to the Michaelis-Menten equation. The measured *K*_m_ for GlcN (black) is more than 10-fold lower than for glucose (gray), in agreement with the observed preference for this substrate. (*F*) Cartoon representation of the three-dimensional structure of SjGlcNK. The N-terminal cap, intermediate subdomain, linker and C-lobe are labeled and colored as in *A*. Arrows indicate the positions of the P-loop, DFD loop and catalytic loop. (*G*) Solid surface representation of SjGlcNK colored according to the electrostatic potential contoured from −8 (red) to 8 (blue) *k*_B_*T*e^−1^ (*k*_B_, Boltzmann’s constant; *T*, temperature (K); e, charge of an electron) highlighting the acidic nature of the active site located in the groove between the intermediate subdomain and the C-lobe.

### SjGlcNK displays a fold similar to mycobacterial maltokinases

The crystal structure of SjGlcNK was solved at 1.98 Å resolution (Table 1) and reveals a typical eukaryotic protein kinase fold, consisting of a N-lobe (formed by a N-terminal cap and an intermediate subdomain) and a C-lobe separated by a linker, with a relative arrangement that results in an overall bean shape (Figure 2*F*). The important conserved catalytic motifs, the DFD loop and the catalytic loop, are located in an acidic groove between the two lobes, whereas the P-loop is in the vicinity of this patch (Figure 2*G*). Structural differences between SjGlcNK and mycobacterial maltokinases are mainly found in the N-terminal cap subdomain. The SjGlcNK N-terminal cap is composed of three long antiparallel β-strands (β**A**-β**B**-β**C**) forming a curved β-sheet enclosing α-helices α**1** and α**2** and by an additional β-strand (β**D**) perpendicular to strand β**B** on its concave surface (Figure S3*A*). In contrast, the N-terminal cap subdomains of mycobacterial maltokinases are 10-14 residues longer than that of SjGlcNK (Figure 2*A*) and display additional secondary structural elements (e.g., β**4**\* and α**B\***), absent in SjGlcNK (Figure S3*A*). These differences could be explained in terms of the different genetic context for both enzymes, since the unique N-terminal cap for the mycobacterial maltokinases is in some cases fused to the C-terminus of the trehalose synthase (Caner et al., 2013; Fraga et al., 2015; PDB entry 5JY7). The intermediate subdomain of SjGlcNK, which is composed by a six-stranded β-sheet (β**F-**β**G-**β**H-**β**J-**β**I-**β**E**) flanked by two α-helical segments (α**3** and α**4**), is almost identical to that of mycobacterial maltokinases, with a root mean square deviation (rmsd) of 1.7 Å for 80 aligned Cα atoms (Figure S3*B*). A seven-residue linker (residues 187-193) connects the intermediate subdomain to the C-terminal lobe (residues 194-438), which is composed of two central 4-helical bundles (α**5**-α**6**-α**10**-α**11** and α**7**-α**8**-α**12**-α**13**) and a small two-stranded β-sheet (β**K**-β**L**), with an additional helix α**9**, also present in MtMak, preceding β**K** and downstream of the catalytic loop. The C-terminal lobe is very similar to the corresponding domain of maltokinases with a rmsd of 1.2 Å upon superposition of 140 Cα atoms (Figure S3*C*).

**Table 1.** Crystallographic data collection and refinement statistics

|  | Apo SjGlcNK<br>(Crystal form A, rendering<br>conformations I and IV)<sup>a</sup> | Apo SjGlcNK<br>(Crystal form B, rendering<br>conformations II, III, IV<br>and V)<sup>a</sup> | Complex of SjGlcNK<br>with ADP, Pi and GLN<br>(Crystal form A, rendering<br>conformations I and VI)<sup>a</sup> |
| --- | --- | --- | --- |
| **Data Collection** |  |  |  |
| X-ray beamline | BL13-XALOC (ALBA) | ID30A-3 (ESRF) | ID30A-3 (ESRF) |
| Space group | $P2_1$ | $P2_1$ | $P2_1$ |
| Cell dimensions | $a = 59.8 \text{ \AA}$<br>$b = 96.1 \text{ \AA}$<br>$c = 80.3 \text{ \AA}$<br>$\beta = 106.7^\circ$ | $a = 58.2 \text{ \AA}$<br>$b = 111.2 \text{ \AA}$<br>$c = 150.6 \text{ \AA}$<br>$\beta = 93.2^\circ$ | $a = 58.8 \text{ \AA}$<br>$b = 97.4 \text{ \AA}$<br>$c = 80.2 \text{ \AA}$<br>$\beta = 107.4^\circ$ |
| Wavelength (Å) | 1.0332 | 0.9677 | 0.9677 |
| Resolution range (Å) | 40.74-1.98 (2.01-1.98) | 47.27-2.69 (2.74-2.69) | 60.18-2.15 (2.19-2.15) |
| Total / Unique reflections | 207,505 / 59,488 (10,538 / 2,936) | 366,841 / 52,817 (19,263 / 2,664) | 325,525 / 46,229 (17,142 / 2,340) |
| Average multiplicity | 3.5 (3.6) | 6.9 (7.2) | 7.0 (7.3) |
| Completeness (%) | 98.1 (97.3) | 98.9 (100.0) | 98.3 (99.2) |
| $R_{\text{meas}}^b$ (%) | 6.7 (106.7) | 22.5 (126.2) | 12.2 (166.0) |
| $CC_{1/2}$ (%) | 99.9 (58.4) | 98.6 (59.0) | 99.8 (55.9) |
| Mean $I/\sigma I$ | 13.2 (1.3) | 9.6 (2.2) | 11.8 (1.3) |
| **Refinement** |  |  |  |
| Resolution range (Å) | 40.7 – 1.98 | 47.3 – 2.69 | 39.9 – 2.15 |
| Unique reflections, work/free | 59,479 / 2,962 | 52,795 / 2,646 | 46,219 / 2,320 |
| R work (%) | 17.6 | 20.6 | 20.7 |
| R free<sup>c</sup> (%) | 21.4 | 24.2 | 24.3 |
| Number of: |  |  |  |
| Amino acid residues | 833 | 1639 | 817 |
| Water molecules | 464 | 256 | 243 |
| Mg<sup>2+</sup> | 7 | 3 | 2 |
| Cl<sup>-</sup> | 4 | 4 | 2 |
| Bis-tris propane | 2 | - | 1 |
| PEG | 3 | - | 3 |
| ADP | - | - | 1 |
| Phosphate ion | - | - | 1 |
| GlcN | - | - | 1 |
| Average B value (Å<sup>2</sup>) |  |  |  |
| Wilson plot | 34.6 | 42.0 | 41.3 |
| Protein | 39.3 (I) / 54.2 (VI) | 44.8 (III) / 50.0 (II) / 46.6 (IV) / 42.7 (V) | 45.7 (I) / 62.9 (VI) |
| Solvent | 49.0 | 38.6 | 50.1 |
| Mg<sup>2+</sup> | 53.5 | 42.9 | 52.6 |
| Cl<sup>-</sup> | 33.7 | 33.4 | 38.3 |
| Bis-tris propane | 96.7 | - | 61.8 |
| PEG | 88.8 | - | 72.5 |
| ADP | - | - | 79.8 |
| Phosphate ion | - | - | 66.8 |
| GlcN | - | - | 73.7 |
| rmsd bond lengths (Å) | 0.006 | 0.002 | 0.002 |
| rmsd bond angles (°) | 0.672 | 0.450 | 0.498 |
| Ramachandran plot<sup>d</sup>,<br>residues in: |  |  |  |
| Favored regions | 801 (97.7%) | 1543 (96.5%) | 776 (97.4%) |
| Additionally allowed | 19 (2.3%) | 55 (3.4%) | 21 (2.6%) |
| Outliers | 0 | 1 (0.1%) | 0 |
| PDB code | 6HWJ | 6HWK | 6HWL |

<sup>a</sup> Numbers in parenthesis correspond to the outermost resolution shell.<sup>b</sup> $R_{\text{meas}}$ is the multiplicity independent R factor (Diederichs and Karplus, 1997).<sup>c</sup> Calculated using 5% of reflections that were not included in the refinement.<sup>d</sup> As defined in MOLPROBITY (Davis et al., 2007).

### Conformational flexibility regulates the catalytic activity of SjGlcNK

SjGlcNK was crystallized in two different monoclinic (*P*2_1_) crystal forms, A and B, which differ considerably in the dimensions of the respective unit cells (Table 1). In consequence, there are two SjGlcNK molecules in the asymmetric unit (AU) of crystal form A, while four monomers are found in the larger AU of crystal form B (Figure S4*A*). The individual structures of the N-terminal cap, the intermediate subdomain, and the C-lobe are highly conserved in the six protomers, across the two crystal forms. In accordance, pairwise superposition of the six structures results in a rmsd of 0.26 - 0.51 Å when the N-terminal cap is aligned, and of 0.30 - 0.50 Å or 0.29 - 0.43 Å when aligning the intermediate subdomain or the C-lobe, respectively. In contrast to the structural conservation of the individual subdomains, the relative positions of the N- and C-lobe are strikingly different in the six molecules (Figure S4*B*). In result, the distance between equivalent atoms in the N- and C-lobe varies from 31.7 Å in the open to 20.6 Å in the closed conformation (Figure 3*A*) and superposition of the C-lobe (residues 194-420) of the six different conformational states results in rmsd values ranging from 0.6 to 5.6 Å for the whole molecule. Principal component (PC) analysis revealed that the six conformers of SjGlcNK are related by an opening-closure movement (PC1), i.e., a hinge-bending motion of the N- and C-lobes of ∼30° amplitude (Figure 3*B*, *C*). In this movement, the linker connecting the two lobes (residues 187-191) acts as a hinge and the catalytic lysine, K133, which sits far from the active site in the open state I (Figure 3*A*, *D*), approaches and establishes contacts with the catalytic D317 in the closed state VI (Figure 3*A*, *E*), likely corresponding to the active conformation of the enzyme.

**Figure 3.**
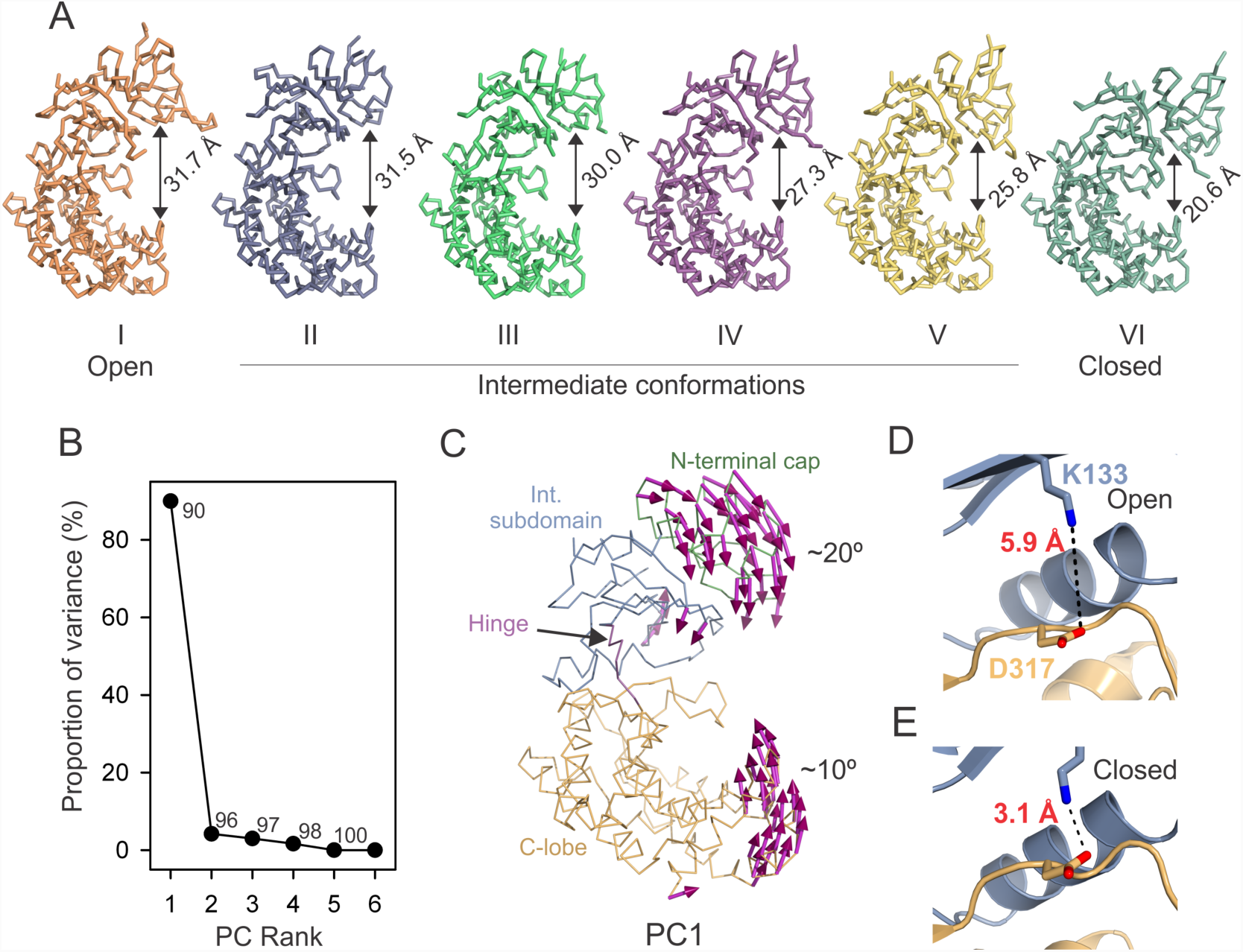
SjGlcNK is highly flexible. (*A*) Ribbon representation of six crystallographic models of SjGlcNK, highlighting their different conformational states. Differences in the relative position of the two lobes (the measured distance between the Cα atoms of A15 and H416 is indicated for each state) indicate conformational flexibility. The structures are labeled with Roman numerals and ordered from left to right, according to decreasing inter-lobe distance. (*B*) Principal component (PC) analysis of subdomain-wide conformational differences between the six crystallographic models. The plot indicates the percentage of the total variance captured by each eigenvector, with numbers indicating the cumulative variance accounted for all preceding eigenvectors. (*C*) Porcupine illustration of the PC1 that relates the six individual SjGlcNK structures. Each subdomain is labeled and colored as in Figure 2*F*. (*D*-*E*) Close-up view of part of the active site of SjGlcNK in the open (*D*) and closed (*E*) conformation. The distance between the catalytic K133 and D317 is indicated.

The quaternary structure and the conformational state of SjGlcNK in solution were elucidated using a combination of small angle X-ray scattering (SAXS) and X-ray crystallography. In solution, SjGlcNK is a monomer (Table S1), adopting an open conformation in the absence of potential substrates (Figure S5*A*). By itself, neither GlcN nor glucose induce apparent conformational changes in SjGlcNK. In contrast, the presence of ATP results in a marked increase in the contribution of closed conformations to the overall scattering. This, together with a decrease in the Guinier radius of gyration (*R*_g_; Figure S5*B*) suggests that upon ATP binding SjGlcNK assumes a more compact and closed conformation. The contribution of a closed conformation to the overall scattering is even slightly larger when SjGlcNK is simultaneously in the presence of ATP and a sugar substrate (Figure S5), suggesting a concerted mechanism where the sugar substrate is phosphorylated in the closed conformation of SjGlcNK. In summary, ATP induces the closure of SjGlcNK and the crystallographic structures reported represent six different conformational snapshots of the enzyme, arising from the molecular flexibility required to carry out its catalytic cycle.

### Caught in the act: Crystal structure of a productive complex of SjGlcNK

In crystals obtained by co-crystallization with a molar excess of ATP and GlcN (Table 1), the substrates could be easily located in the electron density map, at the active site of the enzyme (Figure S6). Interestingly, this structure represents a snapshot of the phosphoryl transfer reaction in which ATP is hydrolyzed into ADP and inorganic phosphate (Pi), with the two hydrolysis products clearly identifiable in the electron density map. The ADP moiety is located in a cleft between the two lobes of SjGlcNK, which adopts a closed conformation (Figure 4*A*), while both Pi and GlcN are found at the C-lobe. Binding of ADP to SjGlcNK is similar to the interaction between the nucleotide and mycobacterial maltokinases (Fraga et al., 2015). The adenine binds to the hinge region, crosslinking both lobes of the enzyme. In particular, the adenine establishes hydrophobic contacts with Y187 and L188 at the hinge, with V131, P164, and V185 at the intermediate subdomain, and with L307 and V316 at the C-lobe. In addition, both the adenine and ribose moieties establish polar contacts with the main chain of A186 and L188 and with the side chain of D193, residues located at the hinge. A major consequence of this extensive network of contacts is the observed ATP-induced closure of SjGlcNK (see above).

**Figure 4.**
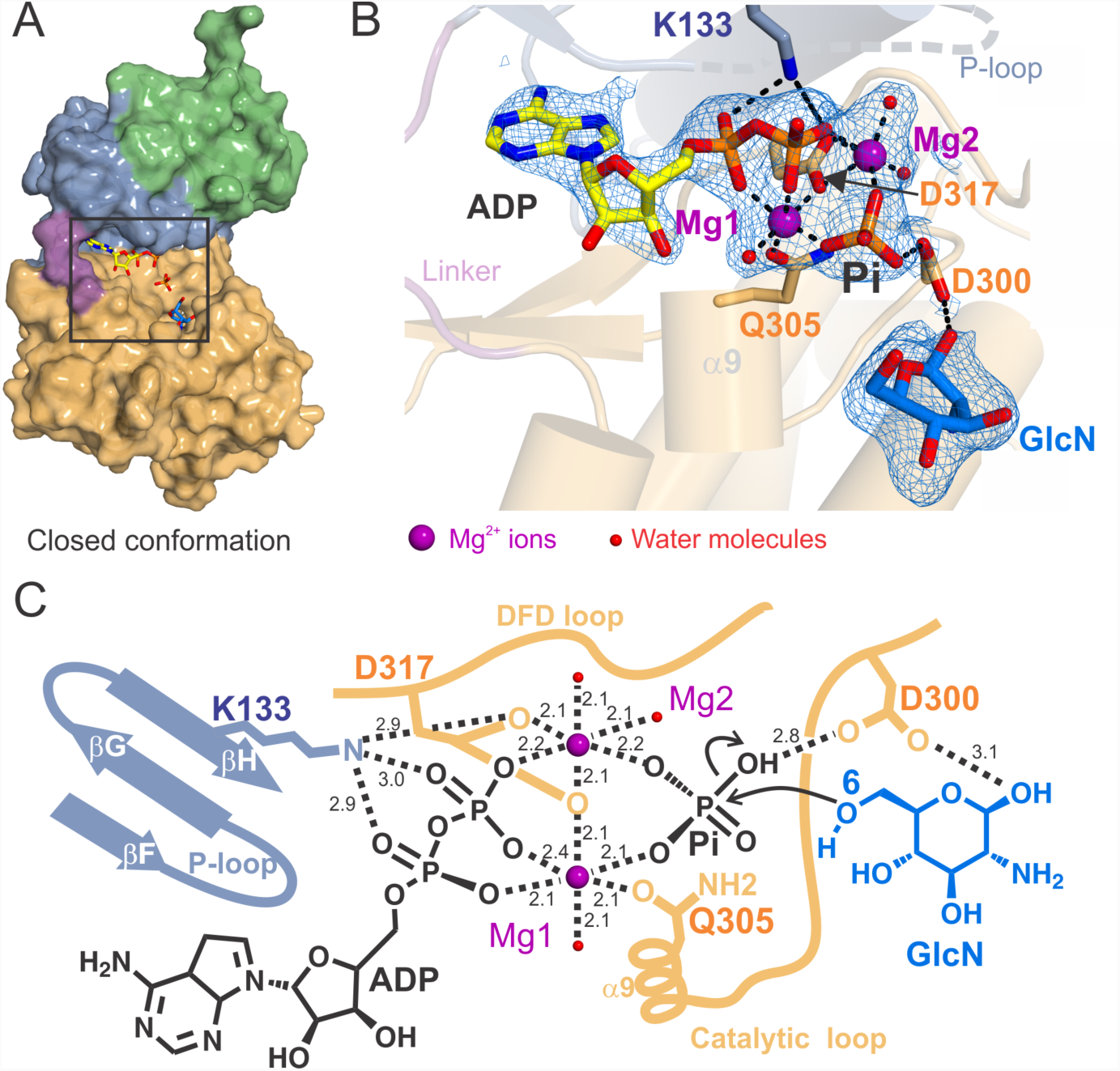
The crystal structure of a productive complex of SjGlcNK unveils the mechanism of phosphate transfer. (*A*) Surface representation of the SjGlcNK-GlcN-ADP-Pi quaternary complex with subdomains colored as in Figure 2*F*. A square highlights the position of the substrates (stick representation) in the active site. (*B*) Close-up view of the enzyme active site in the quaternary complex. The electron density map (2*mF*o-*DF*c contoured at 1.0 σ) around ADP, Pi, GlcN, Mg^2+^ ions and Mg^2+^-coordinating water molecules is shown as a blue mesh. The substrates are shown as sticks with oxygen atoms red, nitrogen blue, phosphorus orange and carbon yellow (ADP) or cyan (GlcN). Metal coordination and hydrogen bonds are indicated by black dashed lines. The protein (intermediate subdomain, linker and C-lobe containing the catalytic helix α9) is depicted in cartoon representation and color coded as in Figure 2*F*. Catalytic residues K133, D300, Q305 and D317 are represented as sticks (carbon atoms colored according to subdomain color scheme). (*C*) Schematic representation of the transition state of GlcN phosphorylation by SjGlcNK. Magnesium ion coordination and polar interactions are shown as black dashed lines, labeled with the respective distances in Å. Magnesium ions and water molecules are shown as magenta and red spheres, respectively.

This crystal structure represents a transient state of GlcN phosphorylation, in which all participating components - ADP, Pi, two magnesium ion cofactors (Mg1 and Mg2), and GlcN - were captured mid-reaction, providing a detailed mechanistic insight into phosphoryl transfer (Figure 4*B*, *C*). In this transition state, the energetically stabilized octahedral coordination of the two magnesium ions drives the breakdown of ATP into ADP and Pi, similar to what is observed for the cAMP-dependent protein kinase (PKA) (Bastidas et al., 2013; Gerlits et al., 2015; Kovalevsky et al., 2012; Zheng et al., 1993). The catalytic residues D317 (in the DFD loop) and Q305 (in the catalytic loop) coordinate the two magnesium ions, stabilizing this transition state. In particular, D317 plays an important role for positioning the Mg1 and Mg2 ions. Curiously, the conformation of the side chain of D317 changes between the nucleotide-free (where it is coordinating the single and more tightly bound Mg1) and the quaternary complex structure (where it coordinates both magnesium ions; Figure S7). The absence of the Mg2 ion in the nucleotide-free structures is not unexpected, since the loss of Mg2 is likely the rate-limiting step of ADP release, as reported in the PKA phosphoryl transfer mechanism (Bastidas et al., 2013).

Although in the quaternary complex K133 establishes hydrogen bonds with the α and β phosphate groups of ADP, this interaction is not key for anchoring the nucleotide to the active site (Carrera et al., 1993). As observed in the SAXS experiments, ATP binding induces more compact and closed conformations of SjGlcNK, bringing K133 closer to the active site (Figure 3*D*, *E*). The positive charge of K133 increases the electrophilicity of the γ-phosphate group of ATP, facilitating its hydrolysis (Matte et al., 1998). In this complex, the leaving phosphate is stabilized by D300 and the two magnesium ions, being perfectly oriented for transfer to position 6 of the bound sugar substrate, GlcN. This unique crystal structure provides experimental evidence of a phosphoryl transfer to GlcN with unprecedented atomic detail on the transition state of the general mechanism of phosphorylation by eukaryotic-like kinases.

### The molecular determinants for substrate specificity in SjGlcNK are highly conserved in Actinobacteria

The functional difference between SjGlcNK and mycobacterial maltokinases, despite their overall structural similarity, is an interesting riddle. Structural superposition of the C-lobes of MtMak and SjGlcNK in complex with maltose and GlcN, respectively, reveals that the sugar binding sites are located in the same position for both enzymes, bordered by the catalytic loop and helices α5, α10 and α12 (Figure 5*A*). Six conserved residues – W195, D300, H302, E409, Y412 and W420 (W222, D322, H324, E430, Y433 and W441 in MtMak) – mediate binding of both sugars (Figure 5*B*). Interestingly, Y370 and R438 that interact with the second glucose moiety of maltose in MtMak are replaced by H348 and L417 in SjGlcNK, which are unable to establish equivalent contacts (Figure 5*C*). In addition, region ^405^QEVR^408^ in helix α12 (^426^KAVY^429^ in MtMak) also plays a part in determining substrate specificity. Noteworthy is the role of Q405, stabilized by polar contacts with R408, in providing specificity to this family of GlcN kinases. The only difference between GlcN and glucose resides at position 2, where the amino group of GlcN is replaced by a hydroxyl. In SjGlcNK, the amino group of GlcN establishes two hydrogen bonds with the side chains of Q405 (K426 in MtMak) and E409 (E430 in MtMak; Figure 5*D*). When Q405 was replaced by an alanine residue in SjGlcNK, its activity remained unchanged towards glucose, but was largely reduced with GlcN (Figure 6*A*). The activity data for SjGlcNK variant Q405A suggest that Q405 is only involved in binding GlcN but not glucose (Figure 6*B*), therefore conferring specificity for the former. Importantly, Q405 and other GlcN-contacting amino acids (D300, H302, R408, E409, Y412 and W420) are highly and exclusively conserved in a large number of uncharacterized aminoglycoside phosphotransferases or putative maltokinases (with lower prediction score) belonging to several representative orders of the phylum *Actinobacteria* (Figure 6*C*). In addition, residues A413, L417, and P418 are also highly conserved. Hence, we propose the consensus sequence “Q-x(2)-RE-x(2)-YA-x(3)-LP-x-W” as a signature for the identification of GlcN kinases, namely those still unclassified in members of the phylum *Actinobacteria*, such as the one described here. Indeed, a BLAST search for this consensus sequence in the non-redundant protein sequences database identified 132 sequences, annotated as putative maltokinases, hypothetical proteins or aminoglycoside phosphotransferases of unknown function, all of them from organisms belonging to the phylum *Actinobacteria* (Figure 6*D*). Interestingly, one of these sequences corresponds to the maltokinase paralogue from *M. smegmatis* (UniProtKB entry: A0A0D6IZ29), MsGlcNK. The substrate specificity of MsGlcNK was also investigated, revealing that it phosphorylates GlcN with a *K*_m_ very similar to that of SjGlcNK, being inactive towards maltose (Figure S8). Thus, the consensus sequence now identified for this unique family of proteins will certainly help to correctly identify and annotate a large number of actinobacterial enzymes, currently with no assigned function or incorrectly annotated as maltokinases.

**Figure 5.**
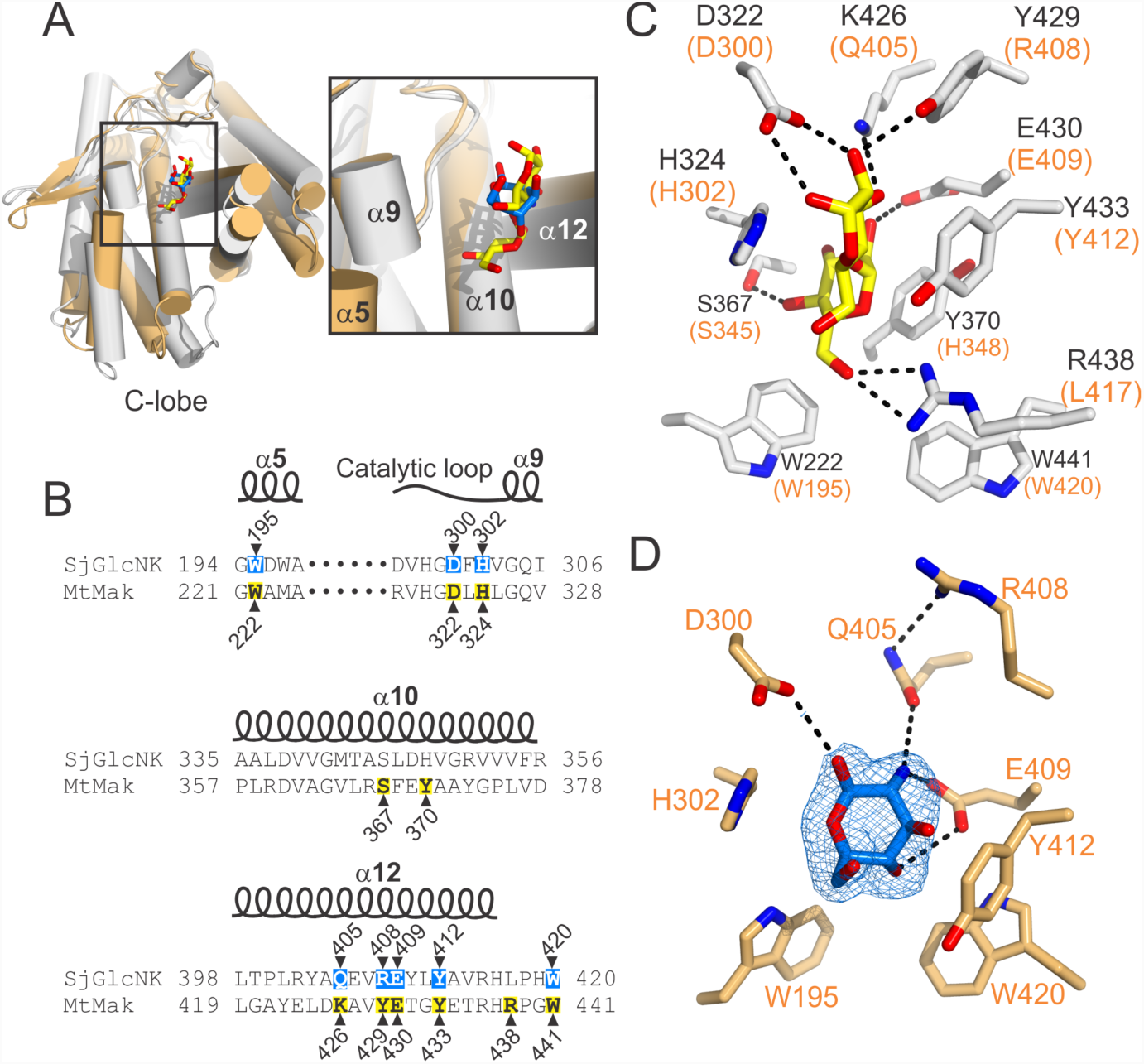
Molecular determinants for substrate specificity in SjGlcNK. (*A*) Cartoon representation of the superposed C-lobes of MtMak (gray, PDB code 4O7P) and SjGlcNK (orange) in complex with maltose (carbon atoms yellow) and GlcN (carbon atoms marine), respectively. The sugar-binding site is highlighted by a square, shown in close-up to the right (key secondary structural elements for the interaction labeled). (*B*) Amino acid sequence alignment of the sugar-binding site of SjGlcNK with that of the mycobacterial maltokinase MtMak (UniProtKB entry O07177). The amino acids interacting with GlcN and maltose are highlighted in cyan and yellow, respectively. Secondary structure elements for SjGlcNK are represented above the alignment. (*C*) Close-up view of the maltose binding-site of MtMak (Li et al., 2014). The amino acids involved in maltose binding and maltose are shown as sticks with oxygen atoms red, nitrogen blue and carbon gray (protein) or yellow (maltose). Residues interacting with maltose are labeled. The equivalent residues in SjGlcNK are indicated in orange brackets. (*D*) Close-up view of the GlcN binding-site of SjGlcNK. The amino acids involved in GlcN binding and GlcN are shown as sticks with oxygen atoms red, nitrogen blue and carbon light orange (protein) or marine (GlcN). Residues interacting with GlcN are labeled. The electron density map (2*mF*o-*DF*c contoured at 1.0 σ) around the bound GlcN molecule is shown as a blue mesh. Dashed black lines represent polar contacts.

**Figure 6.**
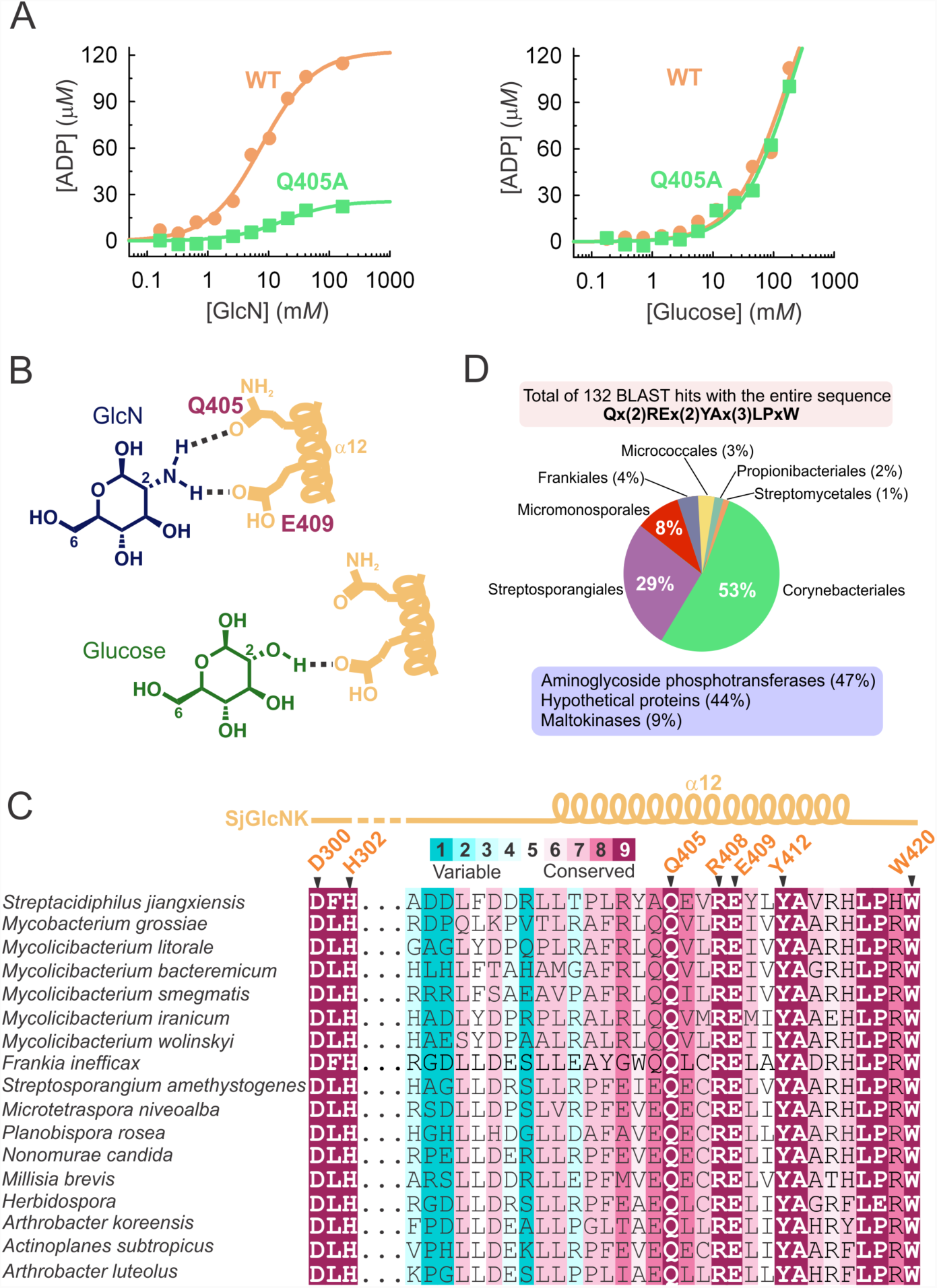
Unveiling the evolutionarily conserved residues for GlcN phosphorylation in *Actinobacteria*. (*A*) Effect of the structure-based amino acid replacement Q405A on the activity of SjGlcNK towards GlcN (left) and glucose (right). The replacement of Q405 by an alanine resulted in a marked decrease of the GlcN-phosphorylating activity of the enzyme, while no effect was observed for glucose, suggesting that Q405 is only involved in GlcN binding. Data were fitted to the Michaelis-Menten equation. (*B*) Schematic representation of the role of Q405 and E409 in GlcN and glucose binding. Both Q405 and E409 are ideally positioned to establish hydrogen bonds with the amino group of GlcN, with Q405 (contact highlighted in blue) being key for GlcN specificity. A binding mode similar to GlcN was assumed for glucose. In contrast to GlcN, glucose establishes no interaction with Q405. (*C*) Multiple amino acid sequence alignment of regions 300-302 and 388-420 of SjGlcNK with the sequences of proposed homologous GlcN kinases from different representative species of *Actinobacteria*. Positions are colored according to evolutionary conservation, calculated from a multiple alignment of 89 unique sequences. The GlcN-contacting residues D300, H302, Q405, R408, E409, Y412 and W420 are highlighted and display strict conservation in these species of *Actinobacteria*. Secondary structure elements for SjGlcNK are represented above the alignment. (*D*) Pie chart indicating the distribution of the 132 hits returned by BLAST when querying the UniProt database with the consensus sequence across the different orders of the phylum *Actinobacteria*. The assigned functions of the hits are given in the light-blue box.

## Discussion

Antibiotic production in *Actinobacteria* is activated through derepression of the regulon of the GntR-family regulator, DasR (Zhu et al., 2014). The regulatory activity of DasR is directly affected by GlcN-6P, which is the central molecule in GlcNAc metabolism (Rigali et al., 2006). To date, two cytosolic pathways leading to GlcN-6P production have been described in *Actinobacteria*: i) deacetylation of GlcNAc-6P, which is formed by phosphorylation of extracellular GlcNAc by the PTS system (Plumbridge and Vimr, 1999; Vogler and Lengeler, 1989; Plumbridge, 2009; Ahangar et al., 2018), and ii) from the glycolytic intermediate fructose-6P by a GlcN-6P synthase (Milewski, 2002; Teplyakov et al., 2002). The uptake of both chitosan-derived GlcN oligosaccharides and monomeric GlcN (from chitin or from peptidoglycan turnover) by the ABC transporter CsnEFG (Viens et al., 2015) opens an alternative mechanism to form GlcN-6P, in which the unique family of GlcN kinases described in this work could be involved. In fact, two other microbial GlcN kinases have been characterized and proposed to catalyze this step in the chitinolytic pathway of their respective organisms. One proteobacterial GlcN-specific kinase, unable to phosphorylate glucose, has been identified in *Vibrio cholerae* (Park et al., 2002) and one archaeal ADP-dependent kinase that phosphorylates glucose and GlcN at similar rates has been identified in *Thermococcus kodakarensis* (Aslam et al., 2018). Sequence homology analyses suggest that the *Vibrio* GlcNK belongs to the ASHKA superfamily of kinases, while the *Thermococcus* kinase is part of the ribokinase-like family. These two non-homologous isofunctional GlcN kinases are now accompanied by the also sequence-unrelated SjGlcNK, a unique GlcN kinase from *Streptacidiphilus jiangxiensis*, an actinomycete that shares the *Streptomycetaceae* family with *Streptomyces*, *Allostreptomyces* and *Kitasatospora*.

Both SjGlcNK and related orthologues were annotated as maltokinases or aminoglycoside phosphotransferases of unknown function. Although the three-dimensional structure of SjGlcNK is very similar to that of mycobacterial maltokinases, it was unable to phosphorylate maltose or aminoglycosides *in vitro*. The molecular determinants for substrate specificity of SjGlcNK were unveiled by the crystal structure of a productive complex with GlcN. The specific interaction between SjGlcNK and GlcN is mediated by a discrete number of highly conserved residues from two short motifs, ^300^D-x-H^302^ in the catalytic loop and ^405^Q-x(2)-RE-x(2)-YA-x(3)-LP-x-W^420^ in helix α12, which we propose as consensus sequences for GlcN phosphorylation in *Actinobacteria*. In accordance, the SjGlcNK homologue from *M. smegmatis*, MsGlcNK, also annotated as maltokinase and displaying the consensus sequence for GlcN binding and processing (Table S2), was shown to phosphorylate GlcN and not maltose (Figure S8). A role for a *Streptomyces coelicolor A3*)*2)* putative saccharide kinase, SCO2662, in the phosphorylation of acquired chitosan-derived GlcN oligosaccharides, has been previously postulated (Viens et al., 2015). Also SCO2662, which belongs to the highly conserved *csnR* gene cluster operon in *Streptomyces* and related genera (e.g., *Kitasatospora*, *Streptosporangium* and *Kribbella*) (Dubeau et al., 2011), displays part of the GlcN-binding consensus sequence, ^255^Q-x(2)-RE-x(2)-AA^263^, although with the tyrosine residue replaced by alanine. Interestingly, a search with the HMMER homologue search algorithm at the Consurf server (Ashkenazy et al., 2016) yielded more than 80 unique sequences of uncharacterized aminoglycoside phosphotransferases or putative maltokinases from organisms of the orders *Corynebacteriales*, *Propionibacteriales*, *Frankiales*, *Streptosporangiales*, *Micromonosporales*, *Micrococcales* and *Streptomycetales* from the phylum *Actinobacteria* (Table S2), in which the identified GlcN-contacting residues are highly conserved. It is worth noting the apparent absence of SjGlcNK homologues not just in other bacterial phyla but also in pathogenic *Actinobacteria*. Indeed, while half of the identified orthologues belong to the *Mycobacteriaceae* family, they were absent from species belonging to the recently emended genera *Mycobacterium* (including *M. tuberculosis* complex and *M. leprae*) and *Mycobacteroides* (including the rapidly growing pathogens *M. abscessus* and *M. chelonae*), which contain the most clinically relevant mycobacterial species (Gupta et al., 2018). Instead, the large majority of mycobacteria with identified orthologues were found to be *Mycolicibacterium* species or strains phylogenetically close to members of this environmental and non-pathogenic genus (Gupta et al., 2018). This was further confirmed by searching the proposed consensus sequence in the non-redundant protein sequences database using BLAST, which resulted in 132 sequences exclusively from the phylum *Actinobacteria*.

While experimental evidence to elucidate the physiological role of these actinobacterial GlcN kinases is still lacking, their genomic context across several genomes strongly suggests a link to secondary metabolism (Figure 1), be it through the activation of the DasR regulon by GlcN-6P or through its putative participation in the biosynthesis of a sugar moiety that may be transferred to the aglycon of a secondary metabolite by its accompanying glycosyltransferase. These observations reinforce the hypothesis that the newly identified kinases have implications in the phosphorylation of acquired extracellular GlcN derived from the hydrolysis of chitosan, i.e., in the incorporation of exogenous GlcN into the bacterial GlcNAc metabolism.

The elucidation of the molecular determinants for GlcN phosphorylation in *Actinobacteria* presented herein will be key for function assignment of a large number of uncharacterized aminoglycoside phosphotransferases, and for reannotation of misannotated bacterial GlcN kinases. In addition, the structural and biochemical characterization of SjGlcNK provides new insights into the role of these unique GlcN kinases as the missing link for the incorporation of environmental GlcN to the metabolism of GlcNAc in bacteria, an important intermediate in peptidoglycan biosynthesis, glycolysis, and secondary metabolite production.

## Materials and methods

### Sequence analyses, genomic context and identification of BGCs

The SjGlcNK amino acid sequence was identified by BLAST searches using the amino acid sequences of characterized mycobacterial maltokinases (EC 2.7.1.175) and retrieved from the PATRIC (http://wwww.patricbrc.org/) database (Genome ID: 235985.3). Paralogue detection and homology between paralogues was determined based on the KEGG SSDB Database (https://www.kegg.jp/kegg/ssdb). For genomic context analyses and identification of BGCs, genome assemblies of *Actinobacteria* containing SjGlcNK homologues were recovered from the NCBI database and analyzed with the antibiotics & Secondary Metabolite Analysis SHell, antiSMASH version 4.2.0 (Blin et al., 2017) using default parameters and inclusion of the ClusterFinder algorithm.

### Strains and culture conditions

*Streptacidiphilus jiangxiensis* 33214 (DSM 45096), obtained from the Deutsche Sammlung von Mikroorganismen und Zellkulturen GmbH (Germany), was cultivated at 28 °C for 6 days in ISP2 agar medium at pH 5.5 (4 g L^−1^ yeast extract, 10 g L^−1^ malt extract, 4 g L^−1^ glucose, 20 g L^−1^ agar). *Mycolicibacterium smegmatis* MC^2^155 (ATCC 700084), obtained from LGC Standards S.L.U. (Spain), was cultivated for 5 days at 30 °C in a glycerol-based agar medium (20 g L^−1^ glycerol, 5 g L^−1^ casamino acids, 1 g L^−1^ fumaric acid, 1 g L^−1^ K_2_HPO_4_, 0.3 g L^−1^ MgSO_4_, 0.02 g L^−1^ FeSO_4_, 2 g L^−1^ Tween 80 with pH 7.0) (Brennan and Ballou, 1967).

### Cloning and site-directed mutagenesis

Chromosomal DNA from *S. jiangxiensis* and from *M. smegmatis* was isolated with the Microbial gDNA Isolation kit (NZYTech). The DNA sequences encoding *S. jiangxiensis* kinase (UniProtKB entry A0A1H7TQR5) and *M. smegmatis* kinase (UniProtKB entry A0A0D6IZ29) were amplified by PCR using KOD Hot-Start DNA polymerase (Novagen) with forward primers 5′-ACATTTCTG<u>CATATG</u>ACCCCGAACTGGT and 5′-TAATAT<u>CATATG</u>ATCGAGCTCGACC and reverse primers 5′-GATC<u>AAGCTT</u>GTCCTTCAGTCTTTCCG and 5′-TAT<u>AAGCTT</u>CGTGCTTGTCCC, respectively (restriction sites underlined). Amplification products were cloned into the NdeI and HindIII sites of pET-30a(+) bacterial expression vector (Novagen), in frame with the vector-encoded C-terminal hexahistidine tag. Single point mutations were introduced by site-directed mutagenesis using the QuikChange method (Stratagene).

### Protein expression and purification

Recombinant *S. jiangxiensis* and *M. smegmatis* proteins were expressed in *Escherichia coli* strain BL21 (DE3). Cells were precultured at 37 °C for 4-5 h in Lysogeny Broth (LB) supplemented with 50 mg L^−1^ kanamycin. Six mL of the preculture were used to inoculate 2-L flasks containing 750 mL LB, 50 mg L^−1^ kanamycin. When cultures reached an OD_600_ of 0.6-0.8 after ∼6 h shaking at 37 °C, protein production was induced by adding 500 μ*M* isopropyl-β-D-thiogalactopyranoside and expression proceeded for ∼16 h at 20 °C. Cells were harvested by centrifugation at 4 °C for 15 min at 3,500 *g* and the resulting pellets were resuspended in 20 m*M* sodium phosphate pH 7.4, 500 m*M* NaCl, 20 m*M* imidazole and frozen at −80 °C. Upon thawing, 1 m*M* PMSF, 1 m*M* DTT, and 1 mg mL^−1^ DNAse were added to the cell suspension and the cells were disrupted by sonication on ice (4 min at 25% amplitude). Cell lysates were centrifuged at 4 °C for 30 min at 39,200 *g* and the supernatant was loaded onto a 5 mL nickel NTA agarose column (Agarose Bead Technologies) equilibrated with 20 m*M* sodium phosphate pH 7.4, 500 m*M* NaCl, 20 m*M* imidazole (Buffer A). Bound protein was eluted with buffer A containing 500 m*M* imidazole. The purity of the eluted fractions was evaluated by SDS-PAGE and those containing recombinant protein were pooled and dialyzed overnight at 4 °C against 20 m*M* Bis-Tris Propane (BTP) pH 7.4, 50 m*M* NaCl, 0.1 m*M* DTT, 0.1 m*M* EDTA. The protein solution was concentrated to 5 mL in a 30 kDa molecular weight cutoff centrifugal ultrafiltration device (Millipore) and loaded onto a Sephacryl S300 HR 26/60 size-exclusion chromatography column (GE Healthcare), with a mobile phase adequate to the downstream use of the protein (see below). Fractions containing pure protein were combined, concentrated by ultrafiltration, flash frozen in liquid N_2_ and stored at −80 °C (for crystallization experiments, the sample was used immediately). Protein concentrations were determined using the extinction coefficient at 280 nm calculated with the ProtParam tool (http://web.expasy.org/protparam/1). Recombinant MvMak was produced and purified as previously described (Fraga et al., 2015).

### Analysis of enzyme activity by thin-layer chromatography

Enzyme activity was assessed in 50 μL mixtures containing 50 m*M* BTP pH 8.0, 10 m*M* MgCl_2_, 20 m*M* phosphate acceptor, 5 m*M* ATP and 2 μg of pure recombinant enzyme at 37 °C. Maltose, maltotriose, isomaltose, trehalose, turanose, leucrose, sucrose, D-mannose, 1-*O*-methylmannose, D-galactose, raffinose, D-xylose, fructose, D-tagatose, D-allose, D-glucose, L-glucose, 3-*O*-methylglucose, 2-deoxyglucose, glucuronic acid, glucosylglycerate, GlcN, GlcNAc, D-mannosamine, *N*-acetyl-D-mannosamine, D-galactosamine, *N*-acetyl-D-galactosamine, *N*-acetylmuramic acid and the aminoglycoside antibiotics kanamycin, streptomycin, gentamicin and hygromycin B were used as potential phosphate acceptors. Reactions were spotted on Silica 60 gel plates (Merck) and developed with one of two solvent systems: acetic acid/ethyl acetate/water/ammonia 25% (6:6:2:1, vol/vol) or ethanol/water (7:3, vol/vol). Sugars were visualized by spraying with α-naphtol/sulfuric acid solution and charring at 120 °C.

### Determination of kinetic parameters

Kinase activity assays were performed with the ADP-Glo^™^ kinase Assay Kit (Promega) (Zegzouti et al., 2009). The reaction mix (containing enzyme, ATP and sugar of interest at different concentrations) was prepared in 100 m*M* Tris pH 7.5, 20 m*M* MgCl_2_, 0.1 mg mL^−1^ BSA. The mix (5 μL) was incubated in an opaque white 384-well assay plate (Corning) at room temperature (RT) for 5 min, followed by addition of the ADP-Glo^™^ reagent (5 μL per well) and incubation for 1 h at RT. Finally, 10 μL of kinase detection reagent were added to each well and the plate was again incubated for 1 h at RT. The luminescence was then recorded using a Synergy 2 plate reader (BioTek) and the data were analyzed with SigmaPlot (Systat Sofware). ATP to ADP standard curves were prepared in the 0.01 μ*M* to 200 μ*M* concentration range.

### Purification of GlcN-6P and NMR analysis

To confirm the identity of the SjGlcNK product a 2 mL reaction mixture (50 m*M* BTP pH 8.0, 20 m*M* MgCl_2_, 20 m*M* GlcN, 10 m*M* ATP and 120 μg SjGlcNK) was incubated overnight at 37 °C and separated by TLC. The mixture was spotted on Silica 60 gel plates (Merck) and developed with a methanol/chloroform/acetic acid/water (4:1:1:1 vol/vol) solvent system. The marginal lanes of the TLC plates were stained to identify the spot corresponding to the SjGlcNK product and the remaining unstained lanes were scrapped in the same region. The product was extracted from the silica gel with ultrapure water, lyophilized and further purified on a Sephadex G10 column (GE Healthcare) with a water flow of 1 mL min^−1^, followed by a second lyophilization step before NMR analysis. All NMR spectra were acquired on an AVANCE III 400 spectrometer from Bruker operating at a central proton frequency of 400.13 MHz equipped with a BBI(F)-z H-X-D (5 mm) probe at 25 ^°^C with presaturation of the water signal.

### Crystallization

All SjGlcNK crystals were obtained by sitting drop vapor diffusion at 20 °C. Crystals belonging to crystal form A grew from drops composed of 1 μL protein solution (45 mg mL^−1^ in 20 m*M* BTP pH 7.4, 50 m*M* NaCl) and 2 μL precipitant (100 m*M* Bis-Tris pH 6.1, 15% (wt/vol) PEG 3350, 0.2 *M* MgCl_2_). Prior to data collection the crystals were transferred to a 1:1 mixture of mineral oil and Parabar 10312 (Hampton Research) and flashed-cooled in liquid N_2_. Crystals of SjGlcNK belonging to crystal form B were obtained from drops composed of 1 μL protein solution (45 mg mL^−1^ in 20 m*M* BTP pH 7.4, 50 m*M* NaCl, pre-incubated for 2 h at 4 °C with 10 m*M* GlcN in the same buffer) and 2 μL crystallization solution (100 m*M* Bis-Tris pH 5.7, 16.5% (wt/vol) PEG 3350, 0.15 *M* MgCl_2_). Prior to data collection the crystals were cryoprotected in crystallization solution supplemented with 20% (vol/vol) glycerol and flashed-cooled in liquid N_2_. Crystals of SjGlcNK in complex with ADP, Pi, and GlcN were obtained from drops composed of 1 μL protein solution (45 mg mL^−1^ in 20 m*M* BTP pH 7.4, 50 m*M* NaCl, pre-incubated for 1 h at 4 °C with 0.2 *M* GlcN, 10 m*M* ATP in the same buffer) and 2 μL precipitant (100 m*M* Bis-Tris pH 6.1, 16% (wt/vol) PEG 3350, 0.15 *M* MgCl_2_). Prior to data collection the crystals were transferred to crystallization solution supplemented with 20% (wt/vol) PEG400 and flashed-cooled in liquid N_2_.

### Data collection and processing

All diffraction data were collected at 100 K. Data from crystal form A (doi:10.15785/SBGRID/614) were collected on the BL13-XALOC beamline (Juanhuix et al., 2014) of the ALBA-CELLS synchrotron (Cerdanyola del Vallès, Spain), and data from crystal form B (doi:10.15785/SBGRID/614) and from the complex with ADP, Pi, and GlcN (doi:10.15785/SBGRID/616) on the ID30A-3 beamline (Theveneau et al., 2013) of the European Synchrotron Radiation Facility (Grenoble, France). Diffraction data were processed with the programs XDS (Kabsch, 2010), Pointless (Evans, 2006), and Aimless (Evans and Murshudov, 2013) as implemented in the autoPROC pipeline (Vonrhein et al., 2011). Crystals belong to the monoclinic space group *P*2_1_ (Table 1) and those of crystal form A and of the complex with ADP, Pi, and GlcN contain two monomers of SjGlcNK in the AU (46% solvent content), while those of crystal form B contain four monomers of SjGlcNK in the AU (51% solvent content).

### Structure solution and refinement

The structure of SjGlcNK was solved by molecular replacement with Phaser (McCoy et al., 2007) as implemented in the MrBUMP pipeline (Keegan and Winn, 2008) from the CCP4 suite (Winn et al., 2011) using data from crystal form A. A marginal solution was found using an ensemble composed by the models of the mycobacterial maltokinases MvMak (PDB entry 4U94; (Fraga et al., 2015)) and MtMak (PDB entry 4O7O; (Li et al., 2014)), but after some rounds of automatic restrained refinement with REFMAC (Murshudov et al., 2011) the *R*_free_ was stuck at 0.52. This model was then subjected to smooth deformation with the morph module of Phenix (Terwilliger et al., 2013) and after 25 morphing cycles (6 Å radius of morphing) there was a substantial improvement in the quality of the electron density maps, accompanied by vastly improved statistics (*R*_work_ = 0.43 and *R*_free_ = 0.47). This initial model was refined with Phenix (Adams et al., 2010), alternating with manual model building with Coot (Emsley et al., 2010). At this stage, only the C-lobe (residues 181-438) could be modeled for the two molecules in the AU. The remaining of molecule A (180 residues) was then built with ARP/wARP (Langer et al., 2008). Finally, the missing segment of molecule B was placed using Phaser and residues 4-180 of molecule A as search model. It then became evident that the two molecules of SjGlcNK in the AU display different conformational states, explaining the difficulties experienced during the phasing stage. The model was completed with alternating cycles of refinement with Phenix (Adams et al., 2010) and manual model building with Coot (Emsley et al., 2010). The final refined model includes residues 1-89, 95-118, and 122-436 of molecule A, and residues 1-19, 33-66, 69-88, 97-115, 123-174, and 178-441 of molecule B, with 97.7% of the main-chain torsion angles in the favored regions of the Ramachandran plot.

The structure of SjGlcNK crystal form B was solved by molecular replacement with Phaser (McCoy et al., 2007) using two fragments (residues 1-188 and 189-433) of molecule A from crystal form A as search models. The resulting solution was refined with Phenix (Adams et al., 2010), alternating with cycles of manual model building with Coot (Emsley et al., 2010). The final refined model (96.5% of the main-chain torsion angles in the favored regions of the Ramachandran plot) comprises residues 4-20, 28-65, 72-89, 93-115, and 121-436 for molecule A, 3-11, 14-19, 35-63, 70-113, and 121-435 for molecule B, 4-20, 27-65, 70-89, 95-117, 122-173 and 177-433 for molecule C, and 3-23, 29-89, 96-117, and 123-434 for molecule D.

Molecular replacement with Phaser (McCoy et al., 2007) using molecules A and B of SjGlcNK crystal form A as search models gave a straightforward solution for the complex with GlcN, ADP and Pi. The model was completed with alternating cycles of refinement with Phenix (Adams et al., 2010) and manual model building with Coot (Emsley et al., 2010). The final refined model (97.4% of the main-chain torsion angles in the favored regions of the Ramachandran plot) comprises residues 1-19, 26-89, 94-117 and 121-433 for molecule A and 4-8, 14-20, 33-63, 72-90, 96-104, 111-116, 121-141 and 143-441 for molecule B. Detailed refinement statistics are given in Table 1. All crystallographic software was supported by SBGrid (Morin et al., 2013).

### SAXS measurements and analysis

SAXS data were collected at BioSAXS beamline BM29 (Pernot et al., 2013) of the European Synchrotron Radiation Facility (Grenoble, France). The SjGlcNK samples were equilibrated in 20 m*M* Tris-HCl pH 8.0, 150 m*M* NaCl, 10 m*M* MgCl_2_, 5 m*M* DTT by size exclusion chromatography (see *Protein expression and purification*, above, for details), concentrated by ultrafiltration on a 10 kDa molecular weight cutoff centrifugal device (Millipore), and centrifuged at 21,500 *g* and 4 °C for 30 min to remove possible aggregates. Mixtures of SjGlcNK (10 mg mL^−1^) with 200 m*M* GlcN, 50 m*M* glucose or 1 m*M* ATP, and the different sugar/nucleotide combinations at these concentrations were prepared and immediately flash frozen in liquid nitrogen. Matching buffers were prepared by adding the adequate amount of ligand(s) to SjGlcNK protein buffer. Prior to data collection, thawed samples were centrifuged at 21,500 *g* and 4 °C for 10 min. All samples and their corresponding buffers were measured consecutively in standard “batch” mode using the automated sample changer, which ensures continuous flow. In order to evaluate the magnitude of interparticle effects, the samples were measured at four concentrations, in the range of 1.25-10.0 mg mL^−1^ or 0.625-5.0 mg mL^−1^, obtained by 2-fold serial dilution of the most concentrated sample. The images (frames) were collected over a scattering vector from 0.0035 to 0.5 Å^−1^ (*q* = (4π sin*θ*)/*λ*, where 2*θ* is the scattering angle). Data were processed and analyzed with the *ATSAS* 2.8 package (Franke et al., 2017). Extrapolation from multiple scattering curves at different concentrations to an infinite dilution and Guinier analysis were done with *PRIMUS*/*qt* (Petoukhov et al., 2012). The *R*_g_ remained constant for all concentrations, within the experimental error. Slight interparticle effects were observed for the two most concentrated samples of apo-SjGlcNK and in presence of GlcN. Hence, to minimize any interparticle effect contribution, the corresponding scattering curves were not used in the extrapolation to infinite dilution. The pair-distance distribution function, *P*(*r*), was calculated with *GNOM* (Svergun, 1992), scattering profiles of atomic structures with *CRYSOL* (Svergun et al., 1995), and the volume fractions for the open (I) and closed (VI) conformations of SjGlcNK with *OLIGOMER* (Konarev et al., 2003).

### Structure and sequence analysis

Subdomain motions were analyzed with DynDom (Hayward Steven and Berendsen Herman J.C., 1998) and evolutionary conservation scores were calculated with Consurf (Ashkenazy et al., 2016). PC analysis was carried out on the collection of experimental structures using the GROMACS utilities (Pronk et al., 2013).

## Data availability

The X-ray diffraction images (http://dx.doi.org/10.15785/SBGRID/614, http://dx.doi.org/10.15785/SBGRID/615 and http://dx.doi.org/10.15785/SBGRID/616) were deposited with the Structural Biology Data Grid (Meyer et al., 2016). Coordinates and structure factors were deposited at the Protein Data Bank (PDB) under accession numbers 6HWJ (SjGlcNK, crystal form A), 6HWK (SjGlcNK, crystal form B) and 6HWL (SjGlcNK-GlcN-ADP-Pi complex). SAXS data were deposited at the Small Angle Scattering Biological Data Bank (SASBDB) (Valentini et al., 2015) under codes SASDEL6, SASDEM6, SASDEN6, SASDEP6, SASDEQ6 and SASDER6. Other data are available from the corresponding authors upon reasonable request.

## Supporting information

## Acknowledgements

We acknowledge the European Synchrotron Radiation Facility (Grenoble, France) for provision of synchrotron radiation facilities and thank their staff for help with data collection. Part of these experiments were performed at beamline BL13-XALOC of ALBA Synchrotron (Cerdanyola del Vallès, Spain), with the collaboration of ALBA staff and CALIPSOplus (Grant 730872) funding. The support of the X-ray Crystallography Scientific Platform of i3S (Porto, Portugal) is also acknowledged. This work was supported by the “Structured program on bioengineered therapies for infectious diseases and tissue regeneration” (Norte-01-0145-FEDER-000012), funded by Norte Portugal Regional Operational Programme (NORTE 2020), under the PORTUGAL 2020 Partnership Agreement, through Fundo Europeu de Desenvolvimento Regional (FEDER) and by FEDER through the COMPETE 2020 - Operacional Programme for Competitiveness and Internationalisation (POCI), Portugal 2020 and by Portuguese funds through FCT - Fundação para a Ciência e a Tecnologia/Ministério da Ciência, Tecnologia e Ensino Superior in the framework of project “Institute for Research and Innovation in Health Sciences” (POCI-01-0145-FEDER-007274) and also by grants UID/NEU/04539/2013 and POCI-01-0145-FEDER-029221. D.N-C acknowledges the European Regional Development Fund (CENTRO-01-0145-FEDER-000012-HealthyAging2020) for a research fellowship and FCT for PhD fellowship SFRH/BD/117777/2016. We thank Dr. Pedro Lamosa from CERMAX, ITQB-NOVA (Oeiras, Portugal) for acquiring and interpreting the NMR data.

## Author contributions

DNC and NE identified the BGCs and cultivated *S. jiangxiensis*; JAM and DNC did the enzymatic assays and substrate specificity determination; JAM and PJBP did the crystallographic studies; JAM did the SAXS analysis; JAM, DNC, SMR, NE and PJBP conceived the experiments and analyzed the data. JAM, SMR, NE and PJBP wrote the manuscript with contributions of all authors. All authors read and approved the final manuscript.

## Conflict of interest

The authors declare no competing financial interests.

