## Supplementary material for "Molecular fingerprints for a novel glucosamine kinase family in *Actinobacteria*"

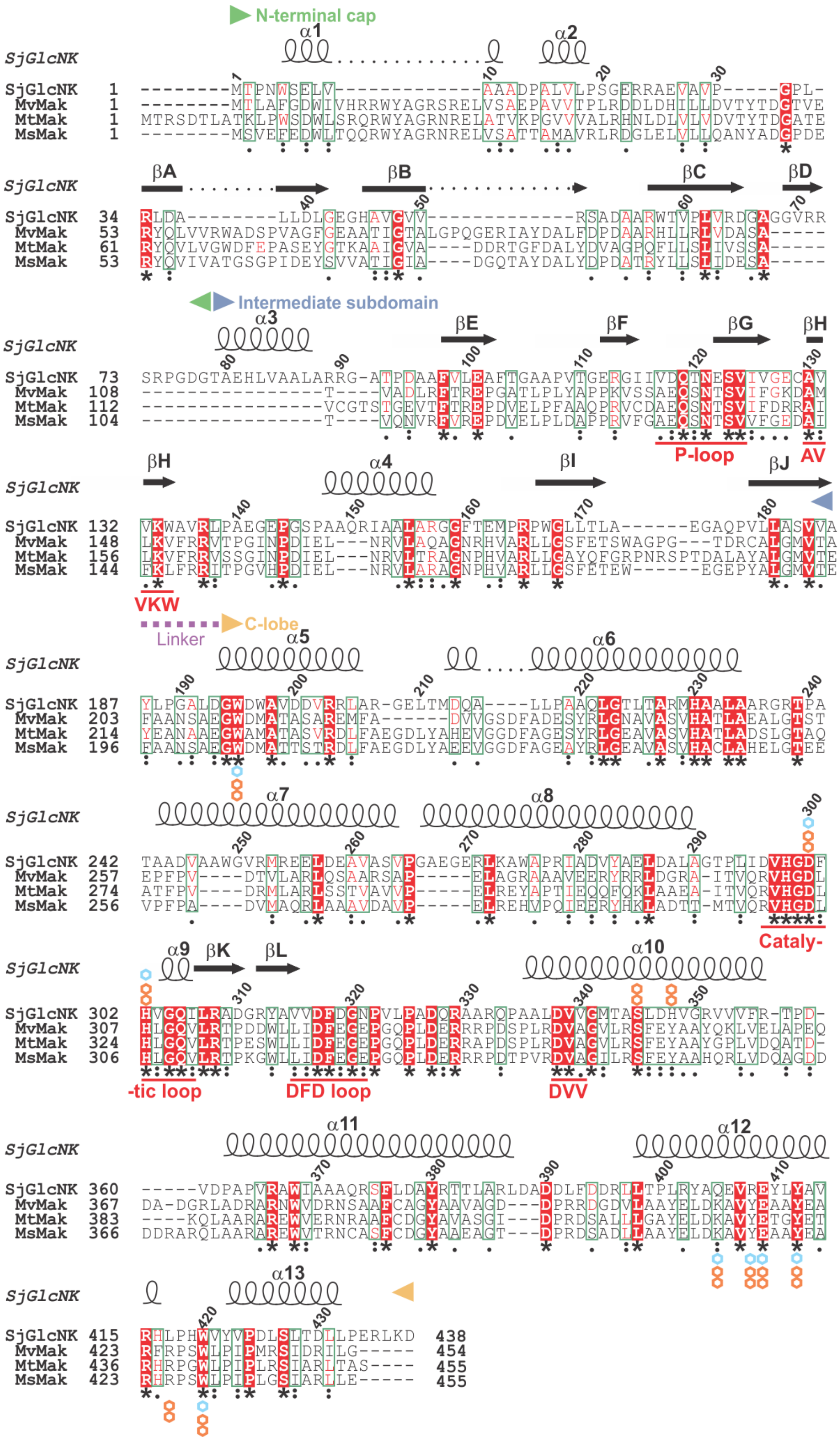

**Figure S1. Multiple amino acid sequence alignment of SjGlcNK with homologous mycobacterial maltokinases.** The amino acid sequence of SjGlcNK from *Streptacidiphilus jiangxiensis* (UniProtKB entry A0A1H7TQR5) was aligned with those of MvMak from *Mycolicibacterium vanbaalenii* (UniProtKB entry A1TH50), MtMak from *Mycobacterium tuberculosis* (UniProtKB entry O07177), and MsMak from *Mycolicibacterium smegmatis* (UniProtKB entry A0R6D9). Strictly conserved alignment positions are shown in inverted type on a red background. Secondary structure elements for SjGlcNK are represented above the alignment. The catalytic and DFD loops, the P-loop, and the DVV and AVVKW motifs are labeled in red. The residues that participate in GlcN and maltose binding are indicated by cyan and orange hexagons, respectively.

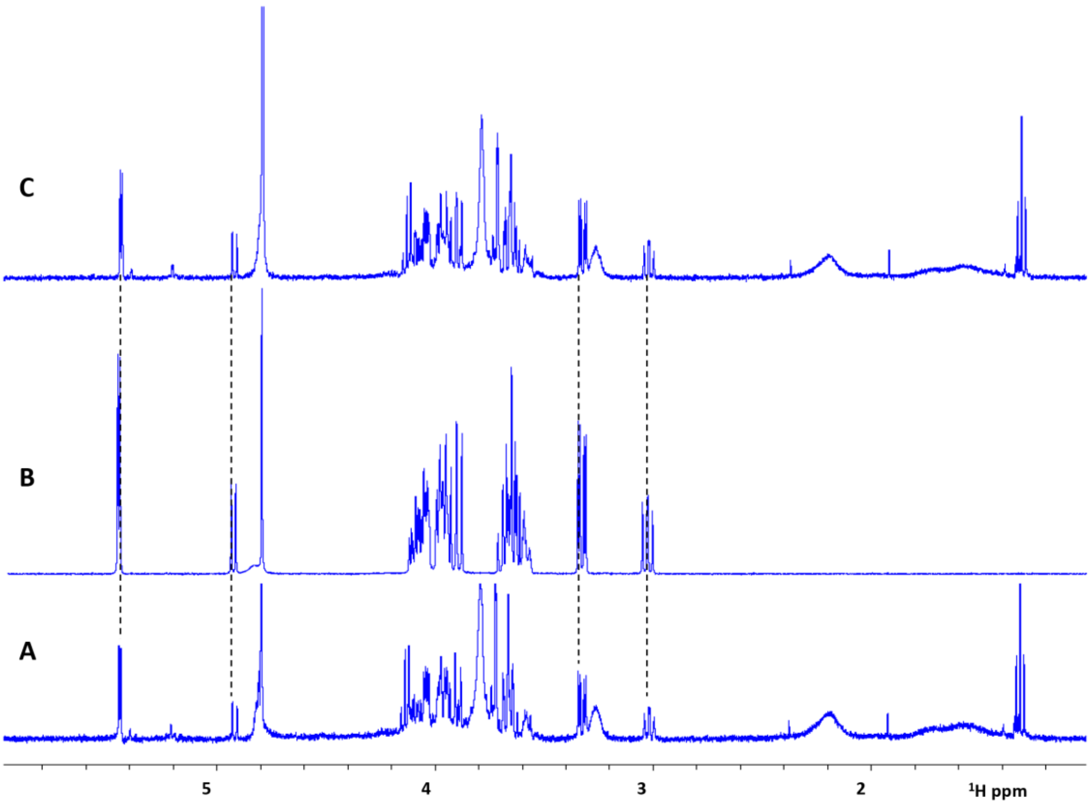


**Figure S2. $^{1}$H-NMR spectra of the enzymatic product of SjGlcNK.** (*A*) Sample from enzymatic reaction mixture purified by thin-layer chromatography; (*B*) GlcN-6P standard (Sigma-Aldrich) in $D_2O$; (*C*) Sample from *A* spiked with an aliquot of standard *B*. The alignment between the signals in samples *A* and *B* is perfect. The spiking of sample *A* with the standard revealed no new signals and a slight increase of those already present in the sample, thus confirming the identity of the sample compound as GlcN-6P.

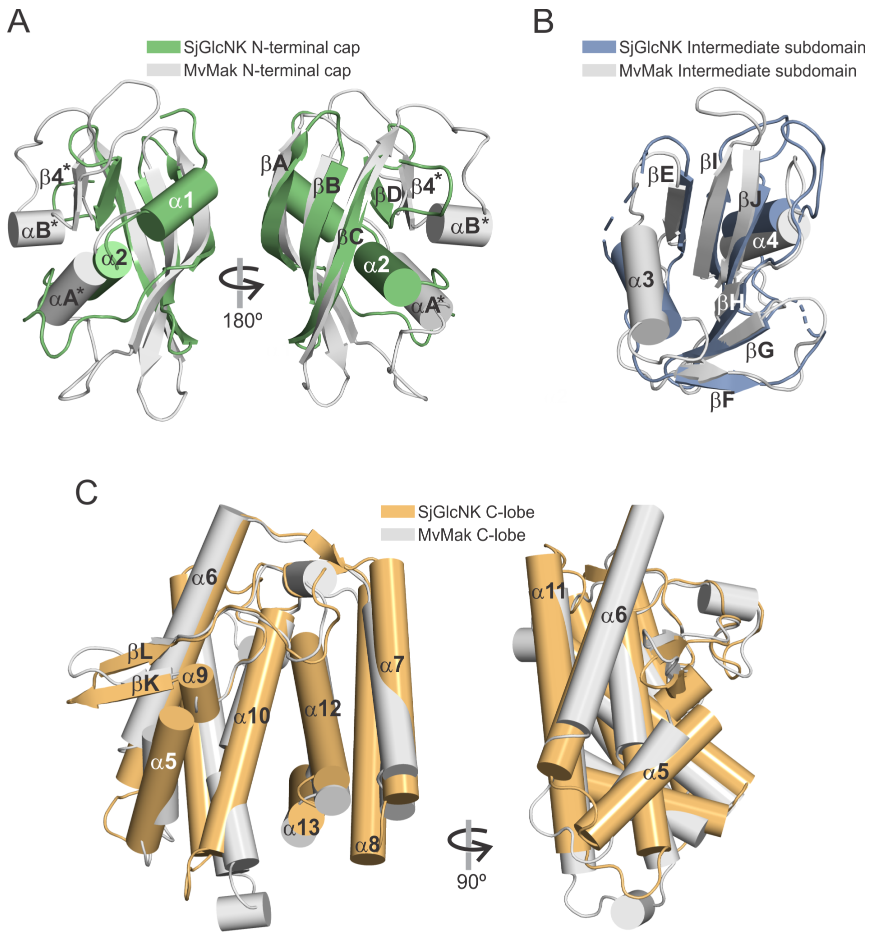


**Figure S3. Overall structure of SjGlcNK and structural comparison with MvMak.** (*A*) Structural superposition of SjGlcNK (colored as in Figure 2*F*) and MvMak (gray; PDB entry 4U94 (Fraga et al., 2015)) N-terminal caps (two views rotated by 180° around *y*), (*B*) intermediate subdomains, and (*C*) C-lobes (two views rotated by 90° around *y*). Secondary structure elements are labeled.

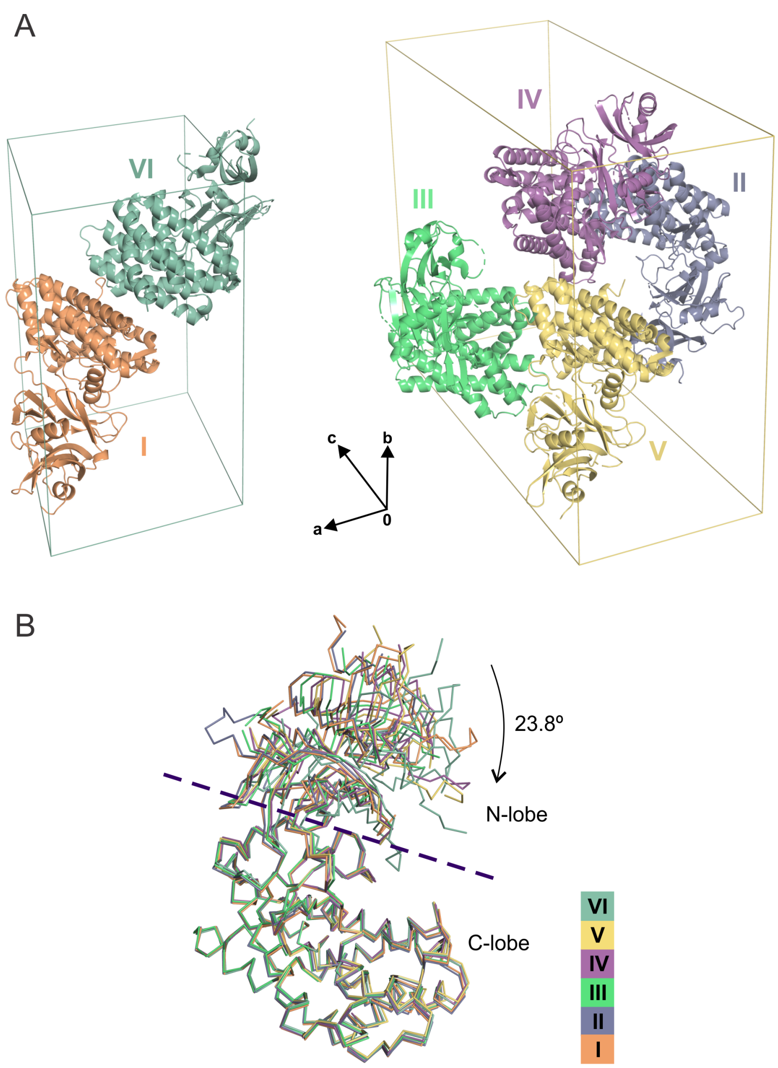


**Figure S4. The two crystallographic forms, A and B, of SjGlcNK comprise six conformational states of the enzyme.** (*A*) Cartoon representation of two (left, crystal form A) and four molecules (right, crystal form B) found in the asymmetric units of the two SjGlcNK crystal forms. The individual structures are labeled with Roman numerals. (*B*) Superposition of the C$\alpha$ traces of the six SjGlcNK monomers (colored as in *A*), by alignment of the C-lobe subdomains. The orientations of the N-lobe in the open and closed conformations are related by a rotation of 23.8° around a hinge axis (dashed line) as determined by DynDom (Hayward Steven and Berendsen Herman J.C., 1998).

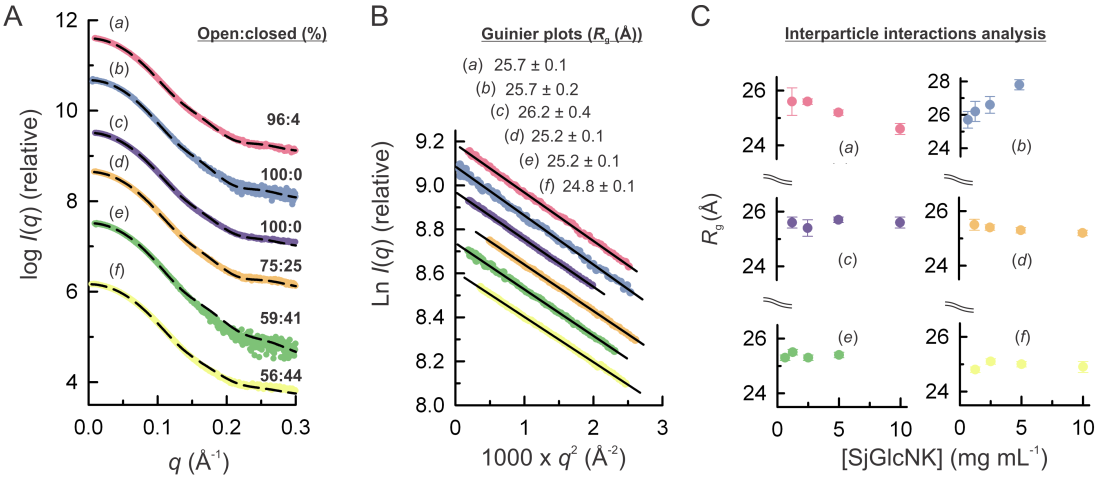


**Figure S5. Ligand-induced conformational transformation of SjGlcNK in solution probed by SAXS.** (*A*) SAXS profiles extrapolated to infinite dilution of SjGlcNK (a) in absence of substrates (red) and in presence of (b) 200 m*M* GlcN (blue), (c) 50 m*M* glucose (violet), (d) 1 m*M* ATP (orange), (e) 200 m*M* GlcN and 1m*M* ATP (green), and (f) 50 m*M* glucose and 1 m*M* ATP (yellow). The curves are offset on the log scale. The scattering calculated for a combination of open and closed conformations (ratio given above the curves), is shown as dashed lines. (*B*) Guinier plots of the scattering data shown in *A*. $R_g$ values (± SD) are indicated for each experiment. (*C*) Plot of Guinier $R_g$ at several concentrations, for the same combinations of protein and ligand(s) as in *A*.

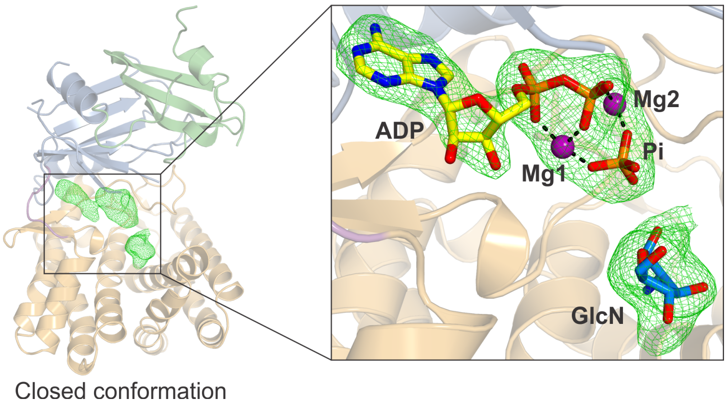


**Figure S6. The crystal structure of the complex between SjGlcNK, GlcN and ATP reveals the transition state of the phosphoryl transfer reaction of ATP to GlcN.** Although a residual, non-interpretable positive electron density was observed near the active site of the closed state (VI in Figure 3*A*) of apo-SjGlcNK (crystal form A), in crystals obtained by co-crystallization with a molar excess of ATP and GlcN (Table 1) the substrates could be easily located in the electron density map. The structure of SjGlcNK (closed conformation) is shown in ribbon representation colored as in Figure 2*F*. The Polder $mF_o$-$DF_c$ map (Liebschner et al., 2017) (contoured at 4$\sigma$) around the ADP, inorganic phosphate (Pi), magnesium ions (Mg1 and Mg2) and GlcN is shown as a green mesh.

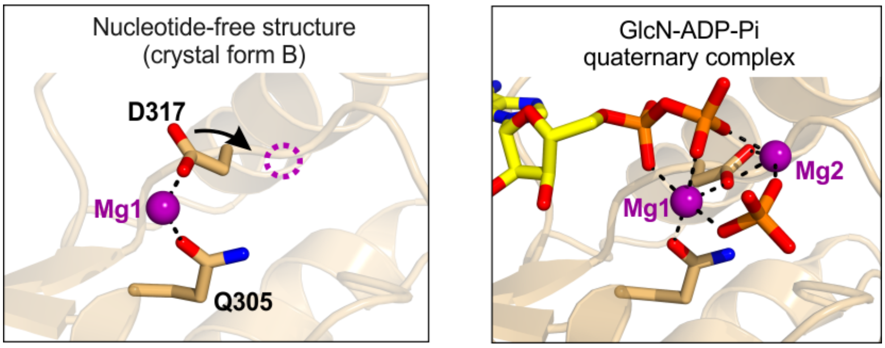


**Figure S7. Ligand binding induces the reorientation of the side chain of D317.** Magnesium binding sites are shown for the nucleotide-free SjGlcNK structure (left) and for the phosphorylation transition state (right) found in the GlcN-ADP-Pi quaternary complex.

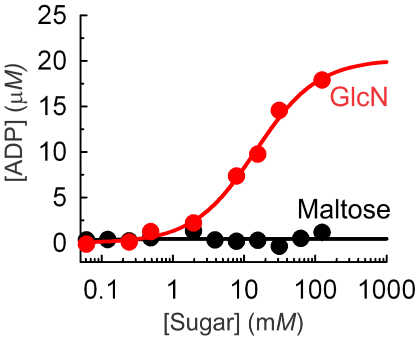


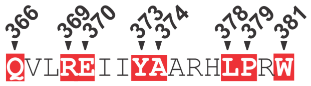


**Figure S8. The putative maltokinase from *M. smegmatis*, MsGlcNK, phosphorylates GlcN.** Similar to SjGlcNK, MsGlcNK displays preference for GlcN over maltose as substrate. The enzymatic activity was measured as ADP release from ATP using the ADP-Glo$^{TM}$ kinase Assay Kit (Promega) (Zegzouti et al., 2009). Kinase reactions were performed in 100 m*M* Tris-HCl pH 7.5, 20 m*M* $MgCl_2$, 0.1 mg $mL^{-1}$ BSA with [MsGlcNK] = 1 μ*M*, [ATP] = 2.5 m*M* and varying concentrations of GlcN (red dots) and maltose (black dots), upon incubation at RT for 5 min. Fitting the data to the Michaelis-Menten equation (red line) resulted in a $K_m$ of 14 ± 2 m*M*, very similar to that of SjGlcNK ($K_m$ = 8 ± 1 m*M*). Part of the amino acid sequence of MsGlcNK is displayed, with the residues matching the proposed consensus sequence Q-x(2)-RE-x(2)-YA-x(3)-LP-x-W for actinobacterial glucosamine kinases highlighted in red.

**Table S1. Small angle X-ray scattering results for SjGlcNK with and without ligands**[a]

(***a***) **Sample details.**

| | |
|---|---|
| Organism | *Streptacidiphilus jiangxiensis* |
| Source | Expressed in *E. coli* BL21 (DE3) |
| UniProtKB entry (residues in construct; C-terminal 6×His-tag residues) | A0A1H7TQR5 (1-451; 439-451) |
| Extinction coefficient [$A_{280}$, 0.1% (*w*/*v*)] | 1.387 |
| $\bar{\upsilon}$ from chemical composition ($cm^3\ g^{-1}$) | 0.742 |
| Particle contrast from sequence and solvent constituents, $\Delta\bar{\rho}$ ($\rho_{protein}$ - $\rho_{solvent}$; $10^{10}\ cm^{-2}$) | 2.75 (12.22 - 9.46) |
| *M* from chemical composition (Da) | 48,249.8 |
| Solvent (solvent blanks taken from SEC flow-through prior to elution of protein) | 20 m*M* Tris-HCl (pH 8.0), 150 m*M* NaCl, 10 m*M* $MgCl_2$, 5 m*M* DTT |

(***b***) **SAXS data collection.**

| | |
|---|---|
| Instrument | ESRF BM29 (Pernot et al., 2013) |
| Detector | Dectris PILATUS 1M |
| Wavelength (Å) | 0.9919 |
| Beam size (μm) | 700 × 700 |
| Camera length (m) | 2.867 |
| *q* measurement range ($Å^{-1}$) | 0.0035-0.5 |
| Absolute scaling method | Comparison with scattering from pure $H_2O$ |
| Normalization | To transmitted intensity by beam-stop counter |
| Monitoring for radiation damage | Data frame-by-frame comparison |
| Exposure time (s) | 10 × 1 |
| Sample configuration | Standard 'batch' mode |
| Sample temperature (ºC) | 10 |

**(*c*) SAXS structural parameters and atomistic modelling.**

| | SjGlcNK | SjGlcNK + 200 m*M* GlcN | SjGlcNK + 1 m*M* ATP | SjGlcNK + 200 m*M* GlcN + 1 m*M* ATP | SjGlcNK + 50 m*M* Glucose | SjGlcNK + 50 m*M* Glucose + 1 m*M* ATP |
|---|---|---|---|---|---|---|
| **Structural parameters** | | | | | | |
| Guinier analysis | | | | | | |
| $I(0)/c$ ($10^{-2}$ cm$^2$ mg$^{-1}$)[b] | 3.280 ± 0.004 | 2.671 ± 0.004 | 3.672 ± 0.002 | 2.637 ± 0.004 | 2.977 ± 0.004 | 3.026 ± 0.002 |
| $R_g$ (Å) | 25.7 ± 0.1 | 25.7 ± 0.1 | 25.2 ± 0.1 | 25.2 ± 0.1 | 26.2 ± 0.4 | 24.8 ± 0.1 |
| $q_{min}$ (Å$^{-1}$) | 0.014 | 0.007 | 0.022 | 0.014 | 0.028 | 0.019 |
| $qR_g$ max | 1.29 | 1.29 | 1.29 | 1.21 | 1.30 | 1.23 |
| Correlation coefficient, $R^2$ | 0.999 | 0.997 | 0.999 | 0.998 | 0.999 | 0.999 |
| *M* from $I(0)/c$ (ratio to predicted)[c] | 45,226 (0.94) | 36,829 (0.76) | 50,659 (1.05) | 36,360 (0.75) | 41,047 (0.85) | 41,724 (0.86) |
| $P(r)$ analysis | | | | | | |
| $I(0)/c$ ($10^{-2}$ cm$^2$ mg$^{-1}$) | 3.283 ± 0.001 | 2.684 ± 0.001 | 3.667 ± 0.002 | 2.640 ± 0.001 | 2.962 ± 0.001 | 3.031 ± 0.001 |
| $R_g$ (Å) | 25.9 ± 0.1 | 26.0 ± 0.2 | 25.2 ± 0.2 | 25.4 ± 0.2 | 26.1 ± 0.2 | 24.9 ± 0.2 |
| $d_{max}$ (Å) | 80 | 80 | 78 | 77 | 80 | 77 |
| $q$ range (Å$^{-1}$) | 0.00913-0.30123 | 0.0090-0.3012 | 0.0223-0.3092 | 0.0137-0.2951 | 0.0279-0.3054 | 0.0190-0.3224 |
| Total estimate from *GNOM* | 0.60 | 0.67 | 0.60 | 0.70 | 0.60 | 0.61 |
| *M* from $I(0)$ (ratio to predicted value) | 45,267 (0.94) | 37,098 (0.77) | 50,562 (1.05) | 36,400 (0.75) | 40,840 (0.85) | 41,791 (0.87) |
| Porod volume (Å$^{-3}$) (ratio $V_P$ / calculated *M*) | 72,000 (1.59) | 78,290 (2.10) | 72,950(1.44) | 75,800 (2.08) | 73,250 (1.79) | 73,100 (1.75) |
| **Atomistic modeling** | | | | | | |
| Contribution (%) of the closed conformation (VI) to the overall scattering as determined *OLIGOMER*[d] | 4 | 0 | 25 | 41 | 0 | 44 |
| SASBDB code[e] | SASDEL6 | SASDEM6 | SASDEN6 | SASDEP6 | SASDEQ6 | SASDER6 |

[a]Description of the accuracy and confidence in the SAXS data and modelling outputs are reported following the *2017 publication guidelines and recommendations for solution small-angle scattering data* (Trewhella et al., 2017). [b]Absolute intensities were determined using water as secondary standard. [c]*M* was calculated as $[N_A\ I(0)/c]/\Delta\rho_M{}^2$, where $I(0)/c$ is the forward scattering normalized against concentration, $\Delta\rho_M = [\rho_{M,prot} - (\rho_{solv}\,\bar{\upsilon})]r_o$ is the scattering contrast per mass, $N_A = 6.023 \times 10^{23}$ mol$^{-1}$ is the Avogadro number, $\rho_{M,prot} = 3.22 \times 10^{23}$ e g$^{-1}$ is the number of electrons per mass of dry protein, $\rho_{solv} = 3.34 \times 10^{23}$ e cm$^{-3}$ is the number of electrons per volume of the aqueous solvent, $\bar{\upsilon}$ is the partial specific volume of the protein and $r_o = 2.8179 \times 10^{-13}$ cm is the scattering length of an electron (Feigin and Svergun, 1987; Mylonas and Svergun, 2007; Orthaber et al., 2000). [d]Form factors for *OLIGOMER* included the calculated intensities for the open (I) and closed (VI) conformations with *CRYSOL*. [e]SASBDB, Small Angle Scattering Biological Data Bank (Valentini et al., 2015).

**Table S2. GlcN kinases proposed as homologues of SjGlcNK in *Actinobacteria* as determined with the ConSurf server (Ashkenazy et al., 2016)**

| Organism | Lineage | Sequence | Protein name | AntiSMASH |
|---|---|---|---|---|
| *Mycobacterium grossiae* | *Actinobacteria Actinobacteria Corynebacteriales Mycobacteriaceae Mycobacterium* | UniRef90_A0A1E8Q961 | Aminoglycoside phosphotransferase | No cluster detected |
| *Mycolicibacterium litorale* | *Actinobacteria Actinobacteria Corynebacteriales Mycobacteriaceae Mycolicibacterium* | UniRef90_A0A1U9PBG4 | Aminoglycoside phosphotransferase | Type I PKS / NRPS cluster |
| *Mycolicibacterium chlorophenolicum* | *Actinobacteria Actinobacteria Corynebacteriales Mycobacteriaceae Mycolicibacterium* | UniRef90_A0A0J6VYP0 | Maltokinase | NRPS cluster |
| *Mycolicibacterium bacteremicum* | *Actinobacteria Actinobacteria Corynebacteriales Mycobacteriaceae Mycolicibacterium* | UniRef90_A0A1W9YXQ7 | Aminoglycoside phosphotransferase | Putative cluster |
| *Mycobacterium* sp. (strain CECT 8779) | *Actinobacteria Actinobacteria Corynebacteriales Mycobacteriaceae Mycobacterium* | UniRef90_UPI000BFECE6E | Aminoglycoside phosphotransferase | Putative cluster |
| *Mycolicibacterium iranicum* | *Actinobacteria Actinobacteria Corynebacteriales Mycobacteriaceae Mycolicibacterium* | UniRef90_A0A1X1WWQ4 | Aminoglycoside phosphotransferase | No cluster detected |
| Uncultured *Mycobacterium* sp. | *Actinobacteria Actinobacteria Corynebacteriales Mycobacteriaceae Mycobacterium* | UniRef90_A0A1Y5PAA8 | Putative 1,4-alpha-glucan branching enzyme | Type I PKS / NRPS cluster |

| | | | | |
|---|---|---|---|---|
| *Mycolicibacterium wolinskyi* | *Actinobacteria*<br>*Actinobacteria*<br>*Corynebacteriales*<br>*Mycobacteriaceae*<br>*Mycolicibacterium* | UniRef90_A0A132PT41 | Aminoglycoside phosphotransferase | Putative cluster |
| *Mycobacterium* sp. (strain 3519A) | *Actinobacteria*<br>*Actinobacteria*<br>*Corynebacteriales*<br>*Mycobacteriaceae*<br>*Mycobacterium* | UniRef90_UPI000C7E100F | Aminoglycoside phosphotransferase | Putative cluster |
| *Mycobacterium* sp. (strain ACS1612) | *Actinobacteria*<br>*Actinobacteria*<br>*Corynebacteriales*<br>*Mycobacteriaceae*<br>*Mycobacterium* | UniRef90_A0A1A1YYA9 | Aminoglycoside phosphotransferase | No cluster detected |
| *Mycobacterium* sp. (strain WY10) | *Actinobacteria*<br>*Actinobacteria*<br>*Corynebacteriales*<br>*Mycobacteriaceae*<br>*Mycobacterium* | UniRef90_A0A1J0UBQ9 | Aminoglycoside phosphotransferase | Type I PKS / NRPS cluster |
| *Mycolicibacterium rufum* | *Actinobacteria*<br>*Actinobacteria*<br>*Corynebacteriales*<br>*Mycobacteriaceae*<br>*Mycolicibacterium* | UniRef90_A0A099CJH1 | Aminoglycoside phosphotransferase | NRPS cluster |
| *Mycolicibacterium rhodesiae* | *Actinobacteria*<br>*Actinobacteria*<br>*Corynebacteriales*<br>*Mycobacteriaceae*<br>*Mycolicibacterium* | UniRef90_A0A1X0J1N8 | Aminoglycoside phosphotransferase | Type I PKS / NRPS cluster |
| *Mycolicibacterium obuense* | *Actinobacteria*<br>*Actinobacteria*<br>*Corynebacteriales*<br>*Mycobacteriaceae*<br>*Mycolicibacterium* | UniRef90_A0A0J6VTN0 | Maltokinase | NRPS cluster |

| | | | | |
|---|---|---|---|---|
| *Mycolicibacterium smegmatis* | *Actinobacteria Actinobacteria Corynebacteriales Mycobacteriaceae Mycolicibacterium* | UniRef90_A0A0D6IZ29 | Trehalose synthase-fused maltokinase | No cluster detected |
| *Mycolicibacterium aromaticivorans* (strain JS19b1) | *Actinobacteria Actinobacteria Corynebacteriales Mycobacteriaceae Mycolicibacterium* | UniRef90_A0A064CHT8 | Aminoglycoside phosphotransferase | Type I PKS / NRPS cluster |
| *Mycobacterium* sp. (strain shizuoka-1) | *Actinobacteria Actinobacteria Corynebacteriales Mycobacteriaceae Mycobacterium* | UniRef90_A0A2C9T723 | Uncharacterized protein | Putative cluster |
| *Mycobacterium* sp. (strain UNC267MFSha1.1M11) | *Actinobacteria Actinobacteria Corynebacteriales Mycobacteriaceae Mycobacterium* | UniRef90_A0A1G4V8L9 | Maltokinase | NRPS cluster |
| *Mycobacterium goodii* | *Actinobacteria Actinobacteria Corynebacteriales Mycobacteriaceae Mycolicibacterium* | UniRef90_A0A0K0X5X8 | Aminoglycoside phosphotransferase | No cluster detected |
| *Mycolicibacterium neoaurum* | *Actinobacteria Actinobacteria Corynebacteriales Mycobacteriaceae Mycolicibacterium* | UniRef90_A0A024QK26 | Trehalose synthase-fused maltokinase | Putative cluster |
| *Mycobacterium* sp. (strain Soil538) | *Actinobacteria Actinobacteria Corynebacteriales Mycobacteriaceae Mycobacterium* | UniRef90_A0A0Q9JII5 | Aminoglycoside phosphotransferase | Putative cluster |
| *Mycobacterium* sp. (strain M26) | *Actinobacteria Actinobacteria Corynebacteriales Mycobacteriaceae* | UniRef90_UPI00073E1503 | Hypothetical protein | No cluster detected |

| | | | | |
|---|---|---|---|---|
| | *Mycobacterium* | | | |
| *Mycolicibacterium agri* | *Actinobacteria* *Actinobacteria* *Corynebacteriales* *Mycobacteriaceae* *Mycolicibacterium* | UniRef90_A0A2A7NDE4 | Aminoglycoside phosphotransferase | Putative cluster |
| *Mycolicibacterium neoaurum* (strain VKM Ac-1815D) | *Actinobacteria* *Actinobacteria* *Corynebacteriales* *Mycobacteriaceae* *Mycolicibacterium* | UniRef90_V5X7B3 | Aminoglycoside phosphotransferase | No cluster detected |
| *Mycobacterium* sp. (strain MS1601) | *Actinobacteria* *Actinobacteria* *Corynebacteriales* *Mycobacteriaceae* *Mycobacterium* | UniRef90_A0A1P8XBH9 | Aminoglycoside phosphotransferase | No cluster detected |
| *Mycolicibacterium vaccae* (strain ATCC 25954) | *Actinobacteria* *Actinobacteria* *Corynebacteriales* *Mycobacteriaceae* *Mycolicibacterium* | UniRef90_K0UL61 | Aminoglycoside phosphotransferase | No cluster detected |
| *Mycobacterium* sp. (strain Root135) | *Actinobacteria* *Actinobacteria* *Corynebacteriales* *Mycobacteriaceae* *Mycobacterium* | UniRef90_A0A0T1W453 | Aminoglycoside phosphotransferase | No cluster detected |
| *Mycolicibacterium brisbanense* | *Actinobacteria* *Actinobacteria* *Corynebacteriales* *Mycobacteriaceae* *Mycolicibacterium* | UniRef90_A0A117I5S0 | Aminoglycoside phosphotransferase | Putative cluster |
| *Mycobacterium* sp. (strain 852013-50091_SCH5140682) | *Actinobacteria* *Actinobacteria* *Corynebacteriales* *Mycobacteriaceae* *Mycobacterium* | UniRef90_A0A1A0XUC1 | Aminoglycoside phosphotransferase | No cluster detected |

| | | | | |
|---|---|---|---|---|
| *Mycobacterium* sp. (strain djl-10) | *Actinobacteria* *Actinobacteria* *Corynebacteriales* *Mycobacteriaceae* *Mycobacterium* | UniRef90_A0A1B1WSU8 | Aminoglycoside phosphotransferase | No cluster detected |
| *Mycobacterium* sp.(strain ITM-2016-00317) | *Actinobacteria* *Actinobacteria* *Corynebacteriales* *Mycobacteriaceae* *Mycolicibacterium* | UniRef90_A0A2S8M089 | Aminoglycoside phosphotransferase | No cluster detected |
| *Mycobacterium* sp. (strain GA-2829) | *Actinobacteria* *Actinobacteria* *Corynebacteriales* *Mycobacteriaceae* *Mycobacterium* | UniRef90_A0A101B6W8 | Aminoglycoside phosphotransferase | No cluster detected |
| *Mycolicibacterium aurum* | *Actinobacteria* *Actinobacteria* *Corynebacteriales* *Mycobacteriaceae* *Mycolicibacterium* | UniRef90_UPI00065E7868 | Aminoglycoside phosphotransferase | NRPS cluster |
| *Mycobacterium dioxanotrophicus* | *Actinobacteria* *Actinobacteria* *Corynebacteriales* *Mycobacteriaceae* *Mycobacterium* | UniRef90_A0A1Y0C060 | Aminoglycoside phosphotransferase | Putative cluster |
| *Mycobacterium* sp. (strain Root265) | *Actinobacteria* *Actinobacteria* *Corynebacteriales* *Mycobacteriaceae* *Mycobacterium* | UniRef90_A0A0Q9B0V8 | Aminoglycoside phosphotransferase | Putative cluster |
| *Mycolicibacterium diernhoferi* | *Actinobacteria* *Actinobacteria* *Corynebacteriales* *Mycobacteriaceae* *Mycolicibacterium* | UniRef90_A0A1Q4H776 | Aminoglycoside phosphotransferase | Putative cluster |
| *Nocardioides lianchengensis* | *Actinobacteria* *Actinobacteria* *Propionibacteriales* | UniRef90_A0A1G6VCL7 | Maltokinase | Putative cluster |

| | | | | |
|---|---|---|---|---|
| | *Nocardioidaceae*<br>*Nocardiodes* | | | |
| *Mycobacterium* sp. (strain NAZ190054) | *Actinobacteria*<br>*Actinobacteria*<br>*Corynebacteriales*<br>*Mycobacteriaceae*<br>*Mycobacterium* | UniRef90_A0A132T811 | Aminoglycoside phosphotransferase | Putative cluster |
| *Frankia* sp. (strain EUN1h) | *Actinobacteria*<br>*Actinobacteria*<br>*Frankiales*<br>*Frankiaceae*<br>*Frankia* | UniRef90_A0A1S1QHA0 | Aminoglycoside phosphotransferase | Putative cluster |
| *Nocardioides psychrotolerans* | *Actinobacteria*<br>*Actinobacteria*<br>*Propionibacteriales*<br>*Nocardioidaceae*<br>*Nocardiodes* | UniRef90_A0A1I3PBM5 | Maltokinase | No cluster detected |
| *Streptosporangium amethystogenes* | *Actinobacteria*<br>*Actinobacteria*<br>*Streptosporangiales*<br>*Streptosporangiaceae*<br>*Streptosporangium* | UniRef90_UPI0006919BC2 | Hypothetical protein | No cluster detected |
| *Frankia inefficax* | *Actinobacteria*<br>*Actinobacteria*<br>*Frankiales*<br>*Frankiaceae*<br>*Frankia* | UniRef90_E3JA12 | Aminoglycoside phosphotransferase | Putative cluster |
| *Frankia* sp. (strain BMG5.36) | *Actinobacteria*<br>*Actinobacteria*<br>*Frankiales*<br>*Frankiaceae*<br>*Frankia* | UniRef90_A0A1S1RDJ1 | Aminoglycoside phosphotransferase | Putative cluster |
| *Microbispora* sp. (strain GMKU363) | *Actinobacteria*<br>*Actinobacteria*<br>*Streptosporangiales*<br>*Streptosporangiaceae*<br>*Microbispora* | UniRef90_UPI0006E25C8F | Hypothetical protein | No cluster detected |
| *Nonomuraea* sp. (strain SBT364) | *Actinobacteria* | UniRef90_UPI00066DC382 | Hypothetical protein | No cluster detected |

| | | | | |
|---|---|---|---|---|
| | *Actinobacteria*<br>*Streptosporangiales*<br>*Streptosporangiaceae*<br>*Nonomuraea* | | | |
| *Frankia* sp. (strain DC12) | *Actinobacteria*<br>*Actinobacteria*<br>*Frankiales*<br>*Frankiaceae*<br>*Frankia* | UniRef90_UPI0005F83CE6 | Aminoglycoside phosphotransferase | Putative cluster |
| *Microbispora* sp. (strain ATCC PTA-5024) | *Actinobacteria*<br>*Actinobacteria*<br>*Streptosporangiales*<br>*Streptosporangiaceae*<br>*Microbispora* | UniRef90_W2EVJ5 | Uncharacterized protein | No cluster detected |
| *Thermoactinospora rubra* | *Actinobacteria*<br>*Actinobacteria*<br>*Streptosporangiales*<br>*Streptosporangiaceae*<br>*Thermoactinospora* | UniRef90_UPI000A1052BC | Hypothetical protein | No cluster detected |
| *Microtetraspora niveoalba* | *Actinobacteria*<br>*Actinobacteria*<br>*Streptosporangiales*<br>*Streptosporangiaceae*<br>*Microtetraspora* | UniRef90_UPI000835E86F | Hypothetical protein | No cluster detected |
| *Streptosporangium canum* | *Actinobacteria*<br>*Actinobacteria*<br>*Streptosporangiales*<br>*Streptosporangiaceae*<br>*Streptosporangium* | UniRef90_A0A1I3I320 | Maltokinase | No cluster detected |
| *Microtetraspora fusca* | *Actinobacteria*<br>*Actinobacteria*<br>*Streptosporangiales*<br>*Streptosporangiaceae*<br>*Microtetraspora* | UniRef90_UPI000836A88C | Hypothetical protein | No cluster detected |
| *Sinosporangium album* | *Actinobacteria*<br>*Actinobacteria*<br>*Streptosporangiales*<br>*Streptosporangiaceae*<br>*Sinosporangium* | UniRef90_A0A1G7VBM6 | Maltokinase | No cluster detected |

| | | | | |
|---|---|---|---|---|
| *Nonomuraea solani* | *Actinobacteria Actinobacteria Streptosporangiales Streptosporangiaceae Nonomuraea* | UniRef90_A0A1H6B4L3 | Maltokinase | No cluster detected |
| *Nonomuraea jiangxiensis* | *Actinobacteria Actinobacteria Streptosporangiales Streptosporangiaceae Nonomuraea* | UniRef90_A0A1G8VHP3 | Maltokinase | No cluster detected |
| *Nonomuraea wenchangensis* | *Actinobacteria Actinobacteria Streptosporangiales Streptosporangiaceae Nonomuraea* | UniRef90_A0A1I0LT16 | Maltokinase | No cluster detected |
| *Nonomuraea indica* | *Actinobacteria Actinobacteria Streptosporangiales Streptosporangiaceae Nonomuraea* | UniRef90_UPI000C7ACCE8 | Aminoglycoside phosphotransferase | No cluster detected |
| *Planobispora rosea* | *Actinobacteria Actinobacteria Streptosporangiales Streptosporangiaceae Planobispora* | UniRef90_UPI00083A6397 | Hypothetical protein | No cluster detected |
| *Micromonospora narathiwatensis* | *Actinobacteria Actinobacteria Micromonosporales Micromonosporaceae Micromonospora* | UniRef90_A0A1A8ZPS6 | Maltokinase | Type I PKS / NRPS cluster |
| *Nonomuraea candida* | *Actinobacteria Actinobacteria Streptosporangiales Streptosporangiaceae Nonomuraea* | UniRef90_UPI0006948DCE | Hypothetical protein | No cluster detected |
| *Streptosporangium subroseum* | *Actinobacteria Actinobacteria Streptosporangiales Streptosporangiaceae* | UniRef90_A0A239G8L3 | Maltokinase | No cluster detected |

| | | | | |
|---|---|---|---|---|
| | *Streptosporangium* | | | |
| *Micromonospora viridifaciens* | *Actinobacteria*<br>*Actinobacteria*<br>*Micromonosporales*<br>*Micromonosporaceae*<br>*Micromonospora* | UniRef90_A0A1C4XCY9 | Maltokinase | No cluster detected |
| *Millisia brevis* | *Actinobacteria*<br>*Actinobacteria*<br>*Corynebacteriales*<br>*Gordoniaceae*<br>*Millisia* | UniRef90_UPI000832ABFF | Hypothetical protein | No cluster detected |
| *Nonomuraea* sp. (strain ATCC 55076) | *Actinobacteria*<br>*Actinobacteria*<br>*Streptosporangiales*<br>*Streptosporangiaceae*<br>*Nonomuraea* | UniRef90_A0A1V0A7D7 | Uncharacterized protein | No cluster detected |
| *Herdidospora cretacea* | *Actinobacteria*<br>*Actinobacteria*<br>*Streptosporangiales*<br>*Streptosporangiaceae*<br>*Herdidospora* | UniRef90_UPI0007744C35 | Hypothetical protein | No cluster detected |
| *Micromonospora* sp. (strain CB01531) | *Actinobacteria*<br>*Actinobacteria*<br>*Micromonosporales*<br>*Micromonosporaceae*<br>*Micromonospora* | UniRef90_A0A1Q4ZS62 | Uncharacterized protein | No cluster detected |
| *Arthrobacter koreensis* | *Actinobacteria*<br>*Actinobacteria*<br>*Micrococcales*<br>*Micrococcaceae*<br>*Arthrobacter* | UniRef90_UPI000B328D64 | Hypothetical protein | No cluster detected |
| *Actinoplanes subtropicus* | *Actinobacteria*<br>*Actinobacteria*<br>*Micromonosporales*<br>*Micromonosporaceae*<br>*Actinoplanes* | UniRef90_UPI0006908AF2 | Hypothetical protein | No cluster detected |
| *Arthrobacter luteolus* | *Actinobacteria*<br>*Actinobacteria* | UniRef90_UPI00082976B6 | Hypothetical protein | No cluster detected |

| | | | | |
|---|---|---|---|---|
| | *Micrococcales*<br>*Micrococcaceae*<br>*Arthrobacter* | | | |
| *Agromyces cerinus* subsp. *cerinus* | *Actinobacteria*<br>*Actinobacteria*<br>*Micrococcales*<br>*Micrococcaceae*<br>*Agromyces* | UniRef90_A0A1N6GDG2 | Maltokinase | No cluster detected |
| *Arthrobacter* sp. (strain Edens01) | *Actinobacteria*<br>*Actinobacteria*<br>*Micrococcales*<br>*Micrococcaceae*<br>*Arthrobacter* | UniRef90_A0A0P7GHY6 | Uncharacterized protein | No cluster detected |
| *Agromyces* sp. (strain Root81) | *Actinobacteria*<br>*Actinobacteria*<br>*Micrococcales*<br>*Micrococcaceae*<br>*Agromyces* | UniRef90_A0A0Q8VX32 | Uncharacterized protein | No cluster detected |
| *Agromyces* sp. (strain CF514) | *Actinobacteria*<br>*Actinobacteria*<br>*Micrococcales*<br>*Micrococcaceae*<br>*Agromyces* | UniRef90_A0A1I6INC3 | Maltokinase | No cluster detected |
| *Agromyces* sp. (strain Leaf222) | *Actinobacteria*<br>*Actinobacteria*<br>*Micrococcales*<br>*Micrococcaceae*<br>*Agromyces* | UniRef90_A0A0Q4HCK4 | Uncharacterized protein | No cluster detected |
| *Mumia flava* | *Actinobacteria*<br>*Actinobacteria*<br>*Propionibacteriales*<br>*Nocardioidaceae*<br>*Mumia* | UniRef90_A0A2M9B6V7 | Maltokinase | No cluster detected |

| | | | | |
|---|---|---|---|---|
| *Sphaerisporangium album* | *Actinobacteria*<br>*Actinobacteria*<br>*Streptosporangiales*<br>*Streptosporangiaceae*<br>*Streptosporangium* | UniRef90_UPI000DE91F25 | Aminoglycoside phosphotransferase | No cluster detected |
| *Jishengella* sp. (strain NA12) | *Actinobacteria*<br>*Actinobacteria*<br>*Micromonosporales*<br>*Micromonosporaceae*<br>*Micromonospora* | UniRef90_A0A2W2DCK5 | Aminoglycoside phosphotransferase | No cluster detected |
| *Homoserinimonas* sp. (strain OAct 916) | *Actinobacteria*<br>*Actinobacteria*<br>*Micrococcales*<br>*Microbacteruaceae*<br>*Homoserinimonas* | UniRef90_UPI000DBEA3B7 | Hypothetical protein | No cluster detected |
| *Actinoplanes lutulentus* | *Actinobacteria*<br>*Actinobacteria*<br>*Micromonosporales*<br>*Micromonosporaceae*<br>*Actinoplanes* | UniRef90_UPI000DB94A95 | Hypothetical protein | No cluster detected |
| *Nonomuraea fuscirosea* | *Actinobacteria*<br>*Actinobacteria*<br>*Streptosporangiales*<br>*Streptosporangiaceae*<br>*Nonomuraea* | UniRef90_A0A2T0MNG9 | Maltokinase | No cluster detected |
| *Mycobacterium* sp. (strain ITM-2016-00316) | *Actinobacteria*<br>*Actinobacteria*<br>*Corynebacteriales*<br>*Mycobacteriaceae*<br>*Mycobacterium* | UniRef_A0A2S8LAU3 | Aminoglycoside phosphotransferase | No cluster detected |
